# Optimized chemogenetic ablation and regeneration of enteric nervous system neurons in zebrafish

**DOI:** 10.1101/2025.07.01.660701

**Authors:** Mujahid Ali Shah, Katherine K. Moran, Helen Rueckert, Abigail V. Sharrock, David F. Ackerley, Jeff S. Mumm, Julia Ganz

## Abstract

The enteric nervous system (ENS) is the intrinsic nervous system of the gut and regulates essential gut functions, including motility, digestion, and immune response, ensuring gut homeostasis. ENS dysfunction or loss is associated with gastrointestinal disorders such as Hirschsprung disease (HSCR). Currently, surgery is the only treatment for HSCR, but it often has lifelong, severe complications. Restoring missing ENS neurons by stimulating endogenous neuronal regeneration presents a promising therapeutic approach for ENS disease. To reveal the cellular-molecular mechanisms regulating neuronal regeneration we study a species capable of robust ENS restoration, the zebrafish. For this, we developed a chemogenetic ablation model in zebrafish using the Gal4/UAS NTR 2.0 system for targeted ENS neuron ablation. Spatially and temporally controlled neuronal death was confirmed by morphological changes, quantification of neuronal loss, and TUNEL assays. We observed an acute immune response that normalizes at 1 day of treatment. Quantification of regenerated neurons demonstrated complete restoration of ENS neuron numbers to control levels by 9 days post treatment, with recovery of gut motility. Among the regenerated neurons, nitrergic, cholinergic and VIPergic subtypes showed full recovery, whereas serotonergic neurons only displayed partial recovery, indicating subtype-specific differences in regenerative capacity and/or timing of cell replacement. Our study establishes a robust platform for dissecting the cellular-molecular mechanisms of ENS regeneration to develop potential treatment approaches for ENS-related diseases.

## Introduction

The enteric nervous system (ENS), the largest subdivision of the peripheral nervous system, consists of an intricate network of neurons and glial cells derived from neural crest progenitors that migrate into the developing gastrointestinal tract (Ganz, 2018; Hao & Young, 2009; Taylor et al., 2016). This complex network plays a pivotal role in regulating essential gastrointestinal functions such as gut motility, nutrient absorption, epithelial secretion, and blood flow (Kuil et al., 2021; Kunze & Furness, 1999; Niesler et al., 2021). Dysfunction or loss of the ENS has been linked to various gastrointestinal disorders such as Hirschsprung disease (HSCR), achalasia, and autism spectrum disorder (Niesler et al., 2021). HSCR, which is characterized by variable degrees ENS neuron loss in the distal gut. HSCR one of the most common congenital anomalies of the ENS, leading to severe intestinal obstruction in neonates, with an estimated incidence of 1 in 5,000 live births (Butler Tjaden & Trainor, 2013). To date, ENS disorders can only be treated symptomatically or by surgical removal of the area with ENS neuron deficits. The current standard treatment for HSCR is surgical resection of the affected gut segment. However, patient outcomes vary widely, and most continue to experience lifelong gastrointestinal defects, highlighting the need for alternative treatments. One of the therapeutic approaches for HSCR is the stimulation of resident stem cells to regenerate missing ENS cells with promising results but varying efficacy depending on the age and the specific gut region (Burns et al., 2016; Joseph et al., 2011; Soret et al., 2020). The successful translation of these approaches requires a comprehensive understanding of the cellular and molecular mechanisms that underlie effective ENS regeneration. Elucidating these mechanisms necessitates the development of a reliable animal model that exhibits robust and reproducible regeneration of ENS neurons following injury.

So far, only a few studies - mostly in mammalian models - have investigated ENS regeneration using chemical, mechanical, or chemical-genetic (chemogenetic) injury. Although these approaches have provided valuable insights, mammalian models show limited and incomplete neuronal regeneration in the ENS (Rueckert & Ganz, 2022; Stavely et al., 2024). In contrast, the zebrafish (*Danio rerio*), a non-mammalian animal model system, is a robust model for studying organ regeneration including various parts of the nervous system such as the brain (Kizil et al., 2012) retina (Hanovice et al., 2019), ENS (El-Nachef & Bronner, 2020; Ohno et al., 2021), but also other organs, such as kidney (Diep et al., 2011), heart (Cao & Poss, 2018), and fin (Romero et al., 2018). Zebrafish also offer numerous experimental advantages, including rapid development, large numbers of offspring, optical transparency that enables live imaging, and a high degree of genetic homology with humans (Howe et al., 2013). Despite these experimental advantages (Kuil et al., 2021; Ganz, 2018), so far, only two studies have studied ENS regeneration in zebrafish using a laser ablation approach (El-Nachef & Bronner, 2020; Ohno et al., 2021). Those studies provided the first evidence that the ENS can regenerate in zebrafish following focal neuronal loss. In particular, Ohno et al., showed that neural crest-derived cells migrate into the ablated region, proliferate, and give rise to new neurons, partially restoring ENS neuron numbers. However, several important issues remain to demonstrate that the ENS fully regenerates in zebrafish: First, the number of regenerated neurons did not reach control levels in the studied time frame. Second, it remains to be shown whether different ENS neuronal subtypes are able to regenerate. Third, it remains unclear whether the regenerated neurons functionally integrated into existing ENS circuits to re-establish normal gut function. In addition, focal laser ablation has several technical limitations, including collateral damage to surrounding tissues, labor-intensive protocols, and restricted ablation of just a few cells within a small gut region, hindering downstream analysis for functional recovery assessment (Ohno et al., 2021).

To overcome these challenges, we leveraged an established approach for spatiotemporally controlled cell ablation: the chemogenetic bacterial nitroreductase (NTR) system. This method utilizes the NTR enzyme, which converts prodrugs, e.g., Metronidazole (MTZ), into cytotoxic agents, thereby selectively killing cells that express NTR (Medico et al., 2001). In zebrafish, this chemogenetic method has been successfully applied in regenerative studies by ablating specific cell populations in heart, pancreas, liver, retina, and other organs (Ariga et al., 2010; Curado et al., 2007, Sharrock et al., 2022; White et al., 2017). However, this method has not yet been tested for the ablation of ENS neurons.

In this study, we took advantage of the precise spatial-temporal control of the *Gal4/UAS* system (Asakawa et al., 2008; Zhang et al., 2019) for the chemogenetic ablation of ENS neurons in postembryonic zebrafish larvae (**Figure 1A**). We additionally employed the most advanced iteration of the nitroreductase technology, NTR 2.0, which provides effective ablation of target cells at ∼10-to 100-fold lower concentrations of MTZ than the original NTR system (Sharrock et al., 2022). Using this approach, we establish a robust experimental paradigm of ENS neuron regeneration by ablating only NTR-expressing neurons through MTZ treatment. Morphological and TUNEL analysis confirmed that the ablated cells underwent cell death with a concomitant acute immune response that normalizes within 1 day post treatment (dpt). By 9 dpt, neuronal recovery was complete, with full recovery of most neuronal subtypes, including nitrergic, cholinergic and VIPergic neurons. Our results demonstrate, for the first time, functional recovery of ENS neuron numbers including restoration of gut motility. These findings pave the way for understanding the cellular and molecular mechanisms of ENS regeneration, a first step for developing new therapeutic approaches for gastrointestinal disorders associated with ENS dysfunction in the future.

**Figure 1.**
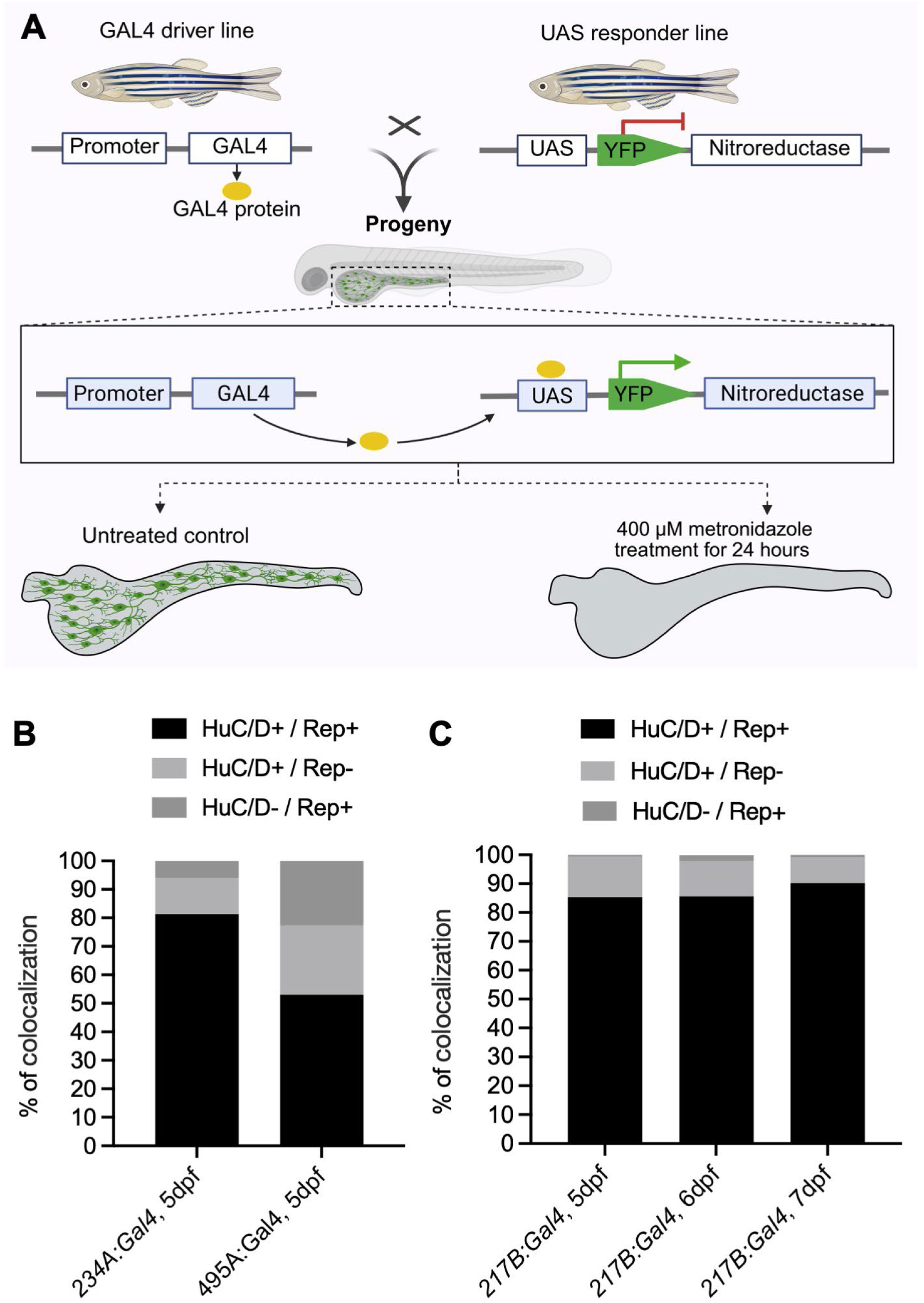
Identification of a Gal4 driver line for efficient ENS neuron ablation using a chemogenetic system. **(A)** Pictorial representation of chemogenetic ablation of ENS neurons. A carrier of the Gal4 driver line, *Tg*(*Gal4)*, which drives expression in ENS neurons was crossed with a carrier of the responder/effector line *Tg(UAS:YFP-NTR 2.0)*, where the Gal4 protein binds to Upstream Activating Sequence (*UAS*) element, initiating expression of both *yellow fluorescence protein* (YFP, reporter protein) and *nitroreductase* (NTR 2.0). Left side: Schematic representation of the untreated control intestine, where enteric neurons co-expressing the *Gal4* and *UAS* transgenes also express both YFP and NTR 2.0 in ENS neurons (green). Right side: Schematic representation of intestine following treatment with 400 µM metronidazole (MTZ). The prodrug MTZ is enzymatically converted by NTR 2.0 into a cytotoxic compound, leading to the selective ablation of YFP-NTR 2.0-expressing enteric neurons. The bar graphs show colocalization percentage of the pan-neuronal marker HuC/D with Reporter (Rep) expression within the gut for Gal4 driver. **(B)** For the *234A:Gal4* line, most Rep+ cells are HuC/D+ at 5 dpf (81.3% HuC/D+/Rep+). The remaining cells consist of 12.7% HuC/D+/ Rep- and 6.0% HuC/D-/Rep*+* (n = 12 larvae per group, 2 experiments). For the *495A:Gal4* line, only 55% cells are HuC/D+/Rep+ at 5 dpf, the remaining cells are 23.5% HuC/D+/ Rep- and 21.2% HuC/D-/Rep*+* (n = 12 larvae per group, 2 experiments). **(C)** For the *217B:Gal4* line, there is a high degree of overlap between Rep and HuC/D expression [(85.3% (5 dpf), 85.7% (6 dpf), and 90.2% (7 dpf)]. The remaining cells consist of HuC/D+/ Rep- [14% (5 dpf) 12.3% (6 dpf) and 9.0% (7 dpf)] and HuC/D-/Rep*+* [0.7% (5 dpf), 2.0% (6 dpf), and 0.8% (7 dpf)], n = 13 (5 dpf), 15 (6 dpf), and 14 (7 dpf) larvae per group, 2 experiments per time point. dpf = days post fertilization.

## Material and methods

### Animals

All experiments were carried out in accordance with animal welfare laws, guidelines, and policies and were approved by the Michigan State University Institutional Animal Care and Use Committee. Zebrafish lines were maintained in a laboratory breeding colony according to established protocols (Westerfield, 2000). Adult zebrafish were bred naturally in system water, and fertilized eggs were collected and transferred to 100 mm petri dishes containing embryo medium. Embryos were allowed to develop at 28.5°C and staged by days post fertilization (dpf) as described earlier (Kimmel et al., 1995; Webb et al., 2009). The zebrafish lines utilized in this study are listed in **Table 1**. Double transgenic larvae were established to co-express YFP and NTR 2.0 in the ENS neurons. For this, the *Tg(5xUAS:GAP-TagYFP-2A-NTR2.0)jh513Tg* reporter/effector line was maintained together with the *Et(2xNRSE-cfos KALT4)gmc617; Tg(5xUAS:GAP-ECFP,he1.1:GAP-ECFP)gmc1913* transgenic fish line. The *Et(2xNRSE:cfos:KalTA4)gmc617* line expresses KalTA4, a zebrafish-optimized Gal4/VP16 fusion protein with neuronal specificity (Xie et al., 2012; Sharrock et al., 2022). We confirmed that the *Tg(SAGFF(LF)217B)* drives reporter expression in ENS neurons (described in the results section), as an incross of *Tg(5xUAS:GAP-TagYFP-2A-NTR2.0)*; *Et(2xNRSE-cfos KALT4)gmc617; Tg(5xUAS:GAP-ECFP,he1.1:GAP-ECFP)gmc1913* did not show YFP expression in ENS neurons. The *Tg(5xUAS:GAP-ECFP,he1.1:GAP-ECFP)gmc1913* line expresses cyan fluorescent protein (CFP), our analysis was focused on YFP expression driven by the Gal4/UAS system. Genotyping of Gal4 driver lines was performed with the following primers for Gal4 (GAL4F: 5’-CGCTACTCTCCCAAAACCAAAAGG-3’; GAL4R: 5’-TCTCTTCCGATGATGATGTCGCAC -3’ using the PCR protocol: 95° x 3 min - [95° x 30s - 60° x 30s - 72° x 30s] x 35 - 72° x 5’ 12° ∞.

**Table 1:** Transgenic lines used in this study.

| Full name | Abbreviation | Source |
| --- | --- | --- |
| *Tg(SAGFF(LF)217B)* | *217B:Gal4* | Asakawa et al., 2008 |
| *Tg(SAGFF(LF)234A)* | *234A:Gal4* | Asakawa et al., 2008 |
| *Tg(gSAIzGFFM495A)* | *495A:Gal4* | Asakawa et al., 2008 |
| *Tg(5xUAS:EGFP)nkuasgfp* | *UAS:GFP* | Kawakami et al., 2010 |
| *Tg(UAS-E1b:NfsB-mCherry)c264* | *UAS:NTR-mCherry* | Davison et al., 2007 |
| *Tg(5xUAS:GAP-TagYFP-2A-NTR2.0)jh513Tg; Et(2xNRSE-cfos KALT4)gmc617; Tg(5xUAS:GAP-ECFP, he1.1:GAP-ECFP)gmc1913* | *UAS:YFP-NTR 2.0\** | Sharrock et al., 2022; Xie et al., 2012 |

\*The *Tg(5xUAS:GAP-TagYFP-2A-NTR2.0)jh513Tg* fish were maintained together with the *Et(2xNRSE-cfos KALT4)gmc617; Tg(5xUAS:GAP-ECFP,he1.1:GAP-ECFP)gmc1913* lines.

### Chemogenetic ablation using Metronidazole

The *217B:Gal4* driver line was crossed with the *UAS:YFP-NTR 2.0* responder line. Offspring from this cross were sorted based on YFP+ cells in the gut into YFP+ and YFP− groups at 5 dpf using a Zeiss Axio ZoomV16 fluorescent stereo microscope. The sorted larvae were then treated at 5 dpf or for the later time points transferred to the nursery until the day of the assay. Prior to MTZ treatment, larvae - except for the 5 dpf groups - were starved for 24 hours to clear the gut of autofluorescent matter. Approximately 50 larvae were transferred to a petri dish (100 mm × 15 mm) containing 40 mL of 400 µM MTZ (M1547 Sigma Aldrich) (Sharrock et al., 2022) in zebrafish system water (diluted from a 5 mM stock solution). The following experimental groups were used: 1) YFP+ MTZ (YFP+ larvae treated with MTZ; 2) YFP+ control (YFP+ larvae, not treated with MTZ); and 3) YFP-MTZ (YFP-larvae, treated with MTZ). Larvae were incubated in the dark for 24 hours to ensure maximal drug exposure as MTZ is light sensitive. After 24 hours, the effects of MTZ treatment on YFP+ cell ablation were assessed using a Zeiss Axio Zoom.V16 fluorescent stereo microscope. After confirmation of cell loss, the MTZ solution was discarded and larvae were returned to the nursery, placed in 2.8-liter tanks (∼40 larvae per tank), and raised until the 21 dpt / 33 dpf.

### ENS neuron regeneration analysis

The regeneration of MTZ-treated YFP+ cells in the gut was assessed at 12 dpf (0 dpt, before ablation) and subsequently at three-day intervals from 13 dpf (1 dpt) to 27 dpf (15 dpt) and compared to YFP+ untreated controls. At each regeneration time point, both control and treated larvae were checked briefly using a Zeiss Axio Zoom.V16 fluorescent stereo microscope, then fixed and immunoassayed (as described below) by confocal microscopy imaging for data acquisition.

### Sample fixation

To ensure optimal imaging conditions, larvae were fasted for 24 hours prior to sampling at the required stages to allow for gut clearance of autofluorescent matter. Then, larvae were euthanized with an overdose of Tricaine Methanesulfonate (MS-222). **For immunolabeling**, euthanized samples were fixed in 4% paraformaldehyde (PFA) prepared in 1x sweet buffer (8% sucrose, 0.2 M CaCl₂, 0.2 M PO₄ buffer, pH 7.3) for 2 hours at room temperature. Then, they were washed subsequently 2 times in 0.5% Phosphate buffer saline-Triton™ X-100 (PBSTX) for 15 mins at room temperature, and stored in PBSTX at 4°C**. For TUNEL assay**, euthanized samples were fixed overnight in 10% Natural buffer saline (NBF) at 4°C, then washed three times in PBS for 5 minutes at room temperature and stored in PBS at 4°C. **For Hybridization Chain Reaction (HCR)**, euthanized larvae were rinsed thoroughly in RNAase free water, and fixed in RNAase free 4% PFA in PBS at 4°C for ∼16 hours, washed three times in PBS for 5 minutes at room temperature, dehydrated in a graded methanol series (50, 70, 90, 95 and 100 % methanol in nuclease free water), and stored in 100% methanol at −20°C for a minimum of 16 hours or up to six months (Ibarra-García-Padilla et al., 2021).

### TUNEL assay

TUNEL assays were performed using the Click-iT™ Plus TUNEL Assay Kits (C10618, Thermo Fisher Scientific) for In Situ Apoptosis Detection. For this, PBS was removed from fixed samples and permeabilization was carried out using Proteinase K (20 µg/mL) for 50 minutes at room temperature, followed by three PBS washes and a deionized water rinse. Then samples were re-fixed in 10% NBF in PBS for 5 minutes at 37°C, followed by three additional PBS washes and another deionized water rinse. To detect DNA strand breaks (i.e., for the positive control), the larvae were incubated in 1 unit of DNase I (Cat. No. 18068-015) diluted into 1× DNase I Reaction Buffer for 30 minutes at room temperature (**Figure S1)**. After that, larvae were preincubated in 150 µL of Terminal deoxynucleotidyl transferase (TdT) buffer for 10 minutes at 37°C, incubated in the 100 µL TdT reaction mixture {94 µL TdT Reaction Buffer, 2 µL 5-ethynyl-2’-deoxyuridine-5’-triphosphate (EdUTP), and 4 µL TdT enzyme} for 30 minutes at 37°C, rinsed in deionized water, washed twice in 3 % bovine serum albumin (BSA) in 0.1% PBS-Triton™ X-100 for 5 minutes each, and one time in PBS. Then the larvae were incubated with the reaction cocktail (90 µL TUNEL Supermix and 10 µL 10× Click-iT Plus TUNEL Reaction Buffer Additive) for 30 minutes at 37°C in the dark, followed by two 3% BSA in PBS washes and two final PBS washes. TUNEL staining was assessed using a Zeiss Axio Zoom.V16 fluorescence stereomicroscope. Immunostaining was subsequently performed under dark conditions, as described below, and larvae were stored in PBS until imaging.

### Immunostaining

Whole-mount immunostaining was performed as previously described (Bandla et al., 2022; Uyttebroek et al., 2010). The following primary antibodies: 5-HT (1:5,000, catalog number 20080), HuC/D (1:1,000, catalog number A-21271), mouse GFP (1:300, catalog number A11120), rabbit GFP (1:1,000, catalog number A11122), nNOS (1:1,000, catalog number GTX133407) and L-plastin, Lcp1 (1:100, catalog number GTX124420) were utilized to perform immunostaining. The primary antibodies were detected using Alexa Fluor 488 and/or 568 secondary antibodies, followed by washes with 0.3% PBSTX. The staining was assessed using Zeiss Axio Zoom.V16 fluorescent stereo microscope and samples were stored in 0.3% PBSTX at 4°C until imaging.

### Hybridization Chain Reaction

The (HCR) kit, probes, *vip* (Accession number: NM_001114553.3) and *slc18a3a* (Accession number: NM_001077550.1), and hairpins were from Molecular Instruments, Inc. (https://www.molecularinstruments.com) and the HCR protocol was performed as previously described (Ibarra-García-Padilla et al., 2021) with minor modifications. The HCR protocol was followed by immunostaining. The fixed samples stored in 100% methanol were rehydrated in graded methanol series, permeabilized with pre-chilled acetone at −20°C for 25 min, treated with 20 μg of Proteinase K for 1 h at room temperature, refixed in 4% PFA in PBS for 20 min, and washed with PBS-Tween before hybridization. For HCR hybridization, samples were pre-hybridized at 37°C, incubated with a probe mixture (4 μL of each 1 μM probe set stock) for 12–16 hours, washed to remove excess probes, and amplified for 12–16 hours, followed by sequential washes (Ibarra-García-Padilla et al., 2021). For antibody staining, samples were blocked in 5% goat serum, incubated with primary anti-mouse GFP (1:300, catalog number A11120) antibodies overnight at 4°C, washed extensively, and incubated with secondary antibodies overnight (Ibarra-García-Padilla et al., 2021). Throughout the protocol, samples were protected from light to prevent photobleaching, ensuring optimal visualization of hybridization and staining results. Afterward, stained samples were stored in PBST at 4°C until imaging.

### Imaging, data acquisition, and quantification

For analysis of colocalization and quantification, intestines were dissected with forceps and mounted on glass slides in 0.3% PBSTX or Fluoromount-G^®^ mounting media (Southern Biotech, catalog number 0100-01) and covered with a coverslip. For the analysis of Gal4-based reporter co-expression with the pan-neuronal marker HuC/D, mounted intestines were imaged on a Zeiss Axio Imager microscope operated through the ZEN software to determine colocalization. Confocal z-stacks images were captured using the 3i spinning disk confocal microscope with a 10, 20, or 60× objectives, operated through Slidebook6 software (Version 6.0.13). For cell quantification, z-stacks were generated from confocal stacks in Slidebook6, which were used for manual cell counts and colocalization analysis. For analysis within gut regions, the images of the intestine were divided into three sections - anterior, middle, and posterior - as previously described (Jianlong Li et al., 2020; Wallace et al., 2005). Cell counts were normalized to 200 µm to account for varying lengths of each of gut regions between larvae and to 1000 µm for the whole gut.

### Analysis of neuron morphology

To analyze morphological changes in neurons after MTZ exposure, control and MTZ-treated larvae were euthanized at the timepoint indicated, their intestines were dissected, mounted on glass slide in PBS and imaged on the 3i spinning disk confocal microscope with a 60× objective, operated through Slidebook6 software (Version 6.0.13).

### Gut anterograde movements analysis

Larvae from control and treated groups were fasted 24 hours prior to analysis, anesthetized with MS-222, and embedded in 0.8% low-melting-point agarose. Videos of the gut region from midgut to posterior were recorded on the Zeiss Axio Zoom.V16 fluorescent stereo microscope with ZEN Imaging Software (ZEN 2.3 blue edition). All time lapse videos were recorded using the same conditions, including exposure time (206s), frame rate (100 frames / 206 sec), and pixels (2752 x 2208). The total number of anterograde, peristaltic wave movements was analyzed from the recorded videos. The Peristaltic Wave Interval (PWI) time was calculated as: PWI = T/N where PWI represents the average time interval (in seconds) between two consecutive anterograde peristaltic movements. The total duration of each video (T, in seconds) and the total number of peristaltic movements observed (N) were recorded.

### Statistical Analysis

For multiple comparisons, one-way and two-way ANOVA were applied to normally distributed data, while the non-parametric Kruskal–Wallis test was used for non-normally distributed data. For pairwise comparisons, unpaired t-tests were used for normally distributed data, and the Mann–Whitney test was used for non-normally distributed data. To evaluate temporal changes across multiple time points in regeneration studies, multiple unpaired t-tests were performed between MTZ-treated and control larvae. For fractional analysis of neuronal colocalization patterns, descriptive statistics were applied. Fold-change analysis was calculated based on the average number of cells in each group. All statistical analyses were performed using GraphPad Prism 9.1 (GraphPad Software, San Diego, CA, USA). A significance threshold of *p* < 0.05 was used to determine statistically significant differences.

## Results

### Identification of Gal4 driver line for specific ablation of ENS neurons

To identify a Gal4 driver line for maximizing the breadth and selectivity of ENS neuron ablation, we crossed three Gal4 gene-trap driver lines (*495A:Gal4; 234A:Gal4; 217B:Gal4*) with a *UAS* reporter line to analyze the extent and specificity of reporter gene expression in the ENS and surrounding tissues. To determine if reporter gene expression within the ENS is primarily restricted to neurons, thereby minimizing off-target ablation, we investigated reporter gene co-expression with the pan-neuronal marker HuC/D in the ENS at 5 dpf (*495A:Gal4xUAS:GFP*, *234A:Gal4xUAS:NTR:DsRed*) and at 5, 6 and 7 dpf (*217B:Gal4xUAS:GFP*) (**Figure 1**).

Our analysis revealed notable differences in the co-expression profiles among the driver lines. The *495A:Gal4* line has previously been reported to show largely gut-restricted expression, but which gut cells express the reporter had not been determined (Kawakami et al., 2010). We found that the intestines of the *495A:Gal4* line displayed only about half HuC/D*+/*Reporter+ cells with many reporter+ cells being HuC/D- and HuC/D+/Reporter-cells (**Figure 1B**). The expression of *234A:Gal4* has been reported to be expressed in the ENS, but also broadly outside the gut (Kawakami et al., 2010; Heanue et al., 2016). In contrast, the *217B:Gal4* had been reported to have reporter expression in the gut with minimal expression outside the gut compared to the *234A:Gal4* line. Consistent with these reports, we found that both *234A:Gal4* and *217B:Gal4* driver lines exhibited higher number of neurons co-expressing HuC/D*+* and Reporter with only a small population of HuC/D-/Reporter+ and HuC/D+/Reporter-cells (**Figure 1B-C**).

Comparatively, both the *234A:Gal4* and *217B:Gal4* lines had sufficient Reporter+ cells in the gut. However, *234A:Gal4* drives extensive reporter gene expression outside the ENS and in ENS progenitor cells (Heanue et al., 2016), which would lead to ablation of additional, non-targeted cells. Therefore, we chose the *217B:Gal4* line to maximize the extent and specificity of ENS neuron ablation (**Figure 1C**).

### Establishing a robust chemogenetic ablation assay for ENS neuron

To establish a robust ENS neuron ablation assay, we used offspring from a cross between the *217B:Gal4* and *UAS:YFP-NTR 2.0* transgenic lines that co-express YFP and NTR 2.0 in cells in the gut (YFP+). To ablate YFP-NTR 2.0 expressing cells, we treated YFP+ larvae with 400 µM MTZ for 24 hours (Sharrock et al., 2022) starting at 5 or 12 dpf. To assess the extent of ENS cell loss, we quantified YFP+ cells at 6 dpf and 13 dpf (1 dpt for MTZ-treated larvae). In untreated controls, no significant difference in YFP+ cells was observed between 5 and 6 dpf, or 12 and 13 dpf. Conversely, MTZ-treated groups showed a near-complete loss of YFP+ cell numbers at 1 dpt compared to 0 dpt (before treatment) and untreated control (**Figure 2A-C**). As 24 h of MTZ treatment showed robust YFP+ cell ablation at both developmental stages, we decided to move forward with MTZ treatment at 12 dpf for further analyses. Embryonic neurogenesis is still very active at 5 dpf (Olden et al., 2008; Roy-Carson et al., 2017). Therefore, ablating neurons at 5 dpf might include processes of developmental neurogenesis rather than of neuronal regeneration, which we aimed to avoid.

**Figure 2.**
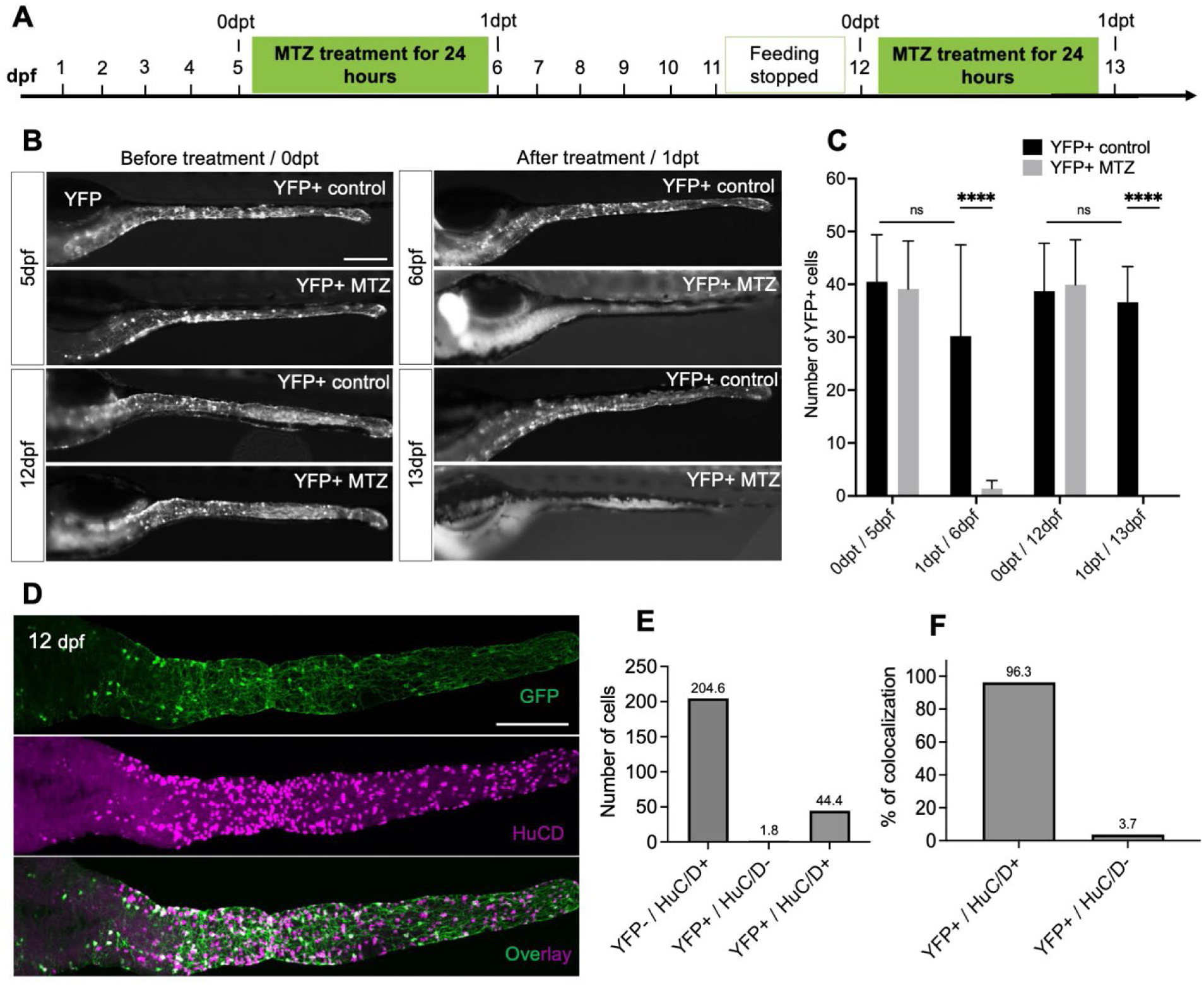
Establishment of a robust ENS neuron ablation assay. **(A)** The timeline of dpf and dpt illustrates the experimental procedure for MTZ treatments initiated at either 5 or 12 dpf. **(B)** Whole-mount side views of live larval guts at the stage indicated. Left panel: YFP+ neurons (white dots) are present before treatment (0 dpt) at 5 or 12 dpf in YFP+ larvae. Right panel: after 24 hours treatment with 400 μM Metronidazole (MTZ), YFP+ cells are completely absent in MTZ-treated fish (YFP+ MTZ) but present in untreated controls (YFP+ control). **(C)** Quantification of YFP+ neurons in control and MTZ-treated groups. Data are shown as mean ± SD (n = 10 larvae per group, 2 experiments); statistical significance \*\*\*\**p* < 0.0001; ns = non-significant. **(D)** At 12 dpf, YFP+ cells [(anti-GFP immunostaining (top, green)] and the pan-neuronal marker HuC/D+ cells (middle, magenta) highly colocalize (bottom, overlay). **(E)** Quantification of GFP+/HuC/D+, GFP+/HuC/D- and GFP-/HuC/D+ cells. Data are shown as mean (n= 10 larvae per group, 2 experiments). **(F)** Percentage of YFP+ cells that are HuC/D+ and HuC/D-cells in **E**. (D) Maximum projection confocal images of dissected intestine at 12 dpf; dpf = days post fertilization. dpf = days post fertilization; dpt = days post treatment; SD = standard deviation; Scale bar = 100 μm.

To confirm that YFP+ cells are neurons at 12 dpf, we analyzed co-expression of YFP and HuC/D in *217B:Gal4*; *UAS:YFP-NTR 2.0* fish. Quantification revealed that, on average, 44.4 YFP+ neurons were observed among a total of 204.6 HuC/D+ neurons. We found that 96.3% of YFP+ cells were HuC/D+ indicating that the vast majority of YFP+ cells are ENS neurons while only 3.7% YFP+ cells are HuC/D-. Therefore, we will hereafter refer to YFP+ cells in these lines as YFP+ neurons (**Figure 2D-F**).

### YFP+ neurons undergo DNA-damage induced cell death after MTZ treatment

To determine the timeline of neuronal ablation, we quantified number of YFP+ neurons at 6, 12 and 24 hpt. We find that YFP+ neurons are essentially absent at 24 hpt indicating that MTZ-induced ablation of YFP+ neurons is completed within 24 hours post-treatment (**Figure S2**). To test if YFP+ neurons undergo cell death after MTZ treatment, we evaluated changes in neuronal morphology and performed TUNEL assays to detect cells undergoing cell death. YFP+ neurons in untreated control larvae displayed normal morphology with no visible signs of morphological changes (**Figure 3A**). In contrast, YFP+ neurons in MTZ-treated samples exhibited significant morphological changes, including axonal and cell body fragmentation and retraction of neuronal processes, consistent with neurons undergoing cell death [**Figure 3**, (Cusack et al., 2013; Ham et al., 2010)]. Additionally, we conducted TUNEL assays combined with immunostaining for YFP to assess DNA damage in control and MTZ-treated fish 6 at hpt. As expected, we found GFP+/TUNEL+ cells in both treated and control groups since apoptotic cell death is a natural ongoing process at 12 dpf (Lee et al., 2023). However, significantly higher numbers of GFP+/TUNEL+ cells were observed in the MTZ-treated group compared to YFP+ untreated control (**Figure 3B, C**). In addition, GFP+/TUNEL+ neurons exhibited clear morphological changes indicating that YFP+ neurons were undergoing cell death (**Figure 3D**). These results indicate that MTZ treatment induces DNA damage which leads to cell death with pronounced structural alterations attending this process, consistent with prior studies using this system for selective cell death (Kulkarni et al., 2018; White & Mumm, 2013).

**Figure 3.**
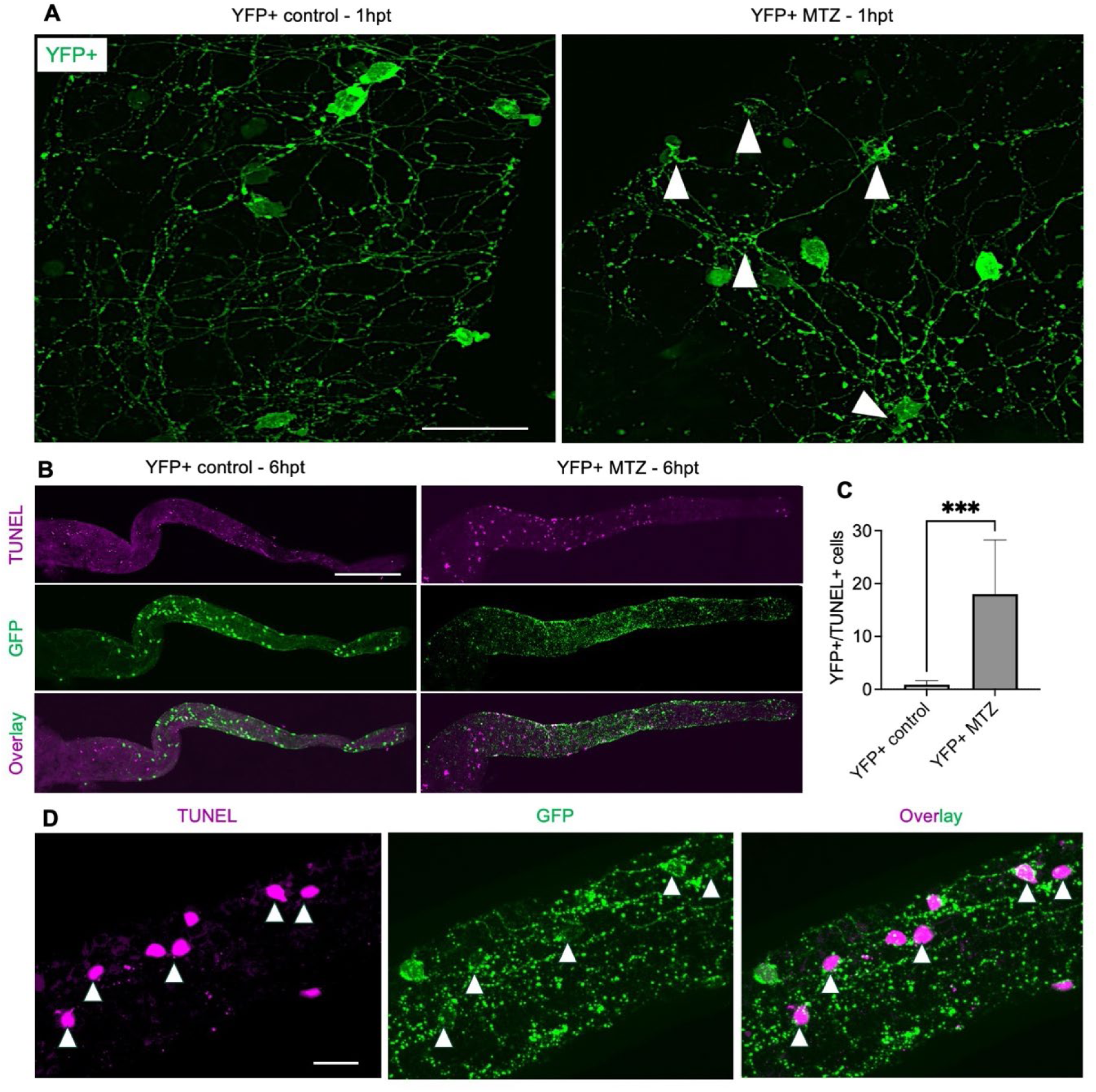
MTZ treatment induces DNA-damage cell death in YFP+ neurons. **(A)** In 12 dpf larvae, YFP+ neurons displays normal neuronal morphology in the untreated control group, while MTZ treated YFP+ neurons show fragmented and retracted neuronal processes at 1 hpt (white arrowheads; n=6 larvae per group, 2 experiments). **(B)** At 6 hpt, there is an increase in TUNEL staining (magenta) in YFP+ neurons (anti-GFP immunostaining) in MTZ treated larvae compared to untreated controls. In YFP+ control group, GFP+ neurons have normal morphology. In contrast, GFP+ neurons in the MTZ-treated group show fragmented morphology indicative of cell death. **(C)** Quantification of GFP+/TUNEL+ cells shows significant increase in YFP+ MTZ-treated larvae compared to YFP+ untreated control. Data are shown as mean ± SD (n= 9 larvae per group, 2 experiments); statistical significance \*\*\**p* < 0.001. **(D)** Magnified images of GFP+/TUNEL+ cells (white arrowhead) in the YFP+ MTZ-treated group. **(A-D)** Maximum projection confocal images of dissected intestines at 1 hpt **(A)** and 6 hpt **(B, D)**. dpf = days post fertilization; hpt = hours post treatment; Scale bars = 20 μm in A; 100 μm in B; 40 μm in D.

### MTZ-induced YFP+ cell ablation elicits acute immune cell activation

Immune cell activation plays a crucial role in neuronal regeneration (Kyritsis et al., 2012; White et al., 2017). To determine whether MTZ-induced cell ablation stimulates an immune response, we analyzed immune cells dynamics using the pan-leukocyte marker L-plastin (Hiroto Shinomiya, 2012). In YFP-MTZ-treated (non-ablated) and YFP+ MTZ-treated (ablated) larvae, we quantified L-plastin+ cell numbers in gut across the time points of neuronal cell death (6 hpt, 12 hpt, and 1 dpt; see **Figure S2**). At 6 hpt, YFP+ MTZ-treated larvae exhibited a significant increase in L-plastin+ cells compared with YFP-MTZ-treated larvae. By 12 hpt and 1 dpt, no significant differences were detected (**Figure 4A, B**). Consistently, fold-change analysis revealed a peak in L-plastin+ cell numbers at 6 hpt, which steadily declined back to normal levels by 1 dpt (**Figure 4C**). These findings indicate that MTZ-induced ablation of YFP+ cells trigger a transient immune response in the gut, characterized by an acute accumulation of L-plastin+ cells that resolves within 24 hours.

**Figure 4.**
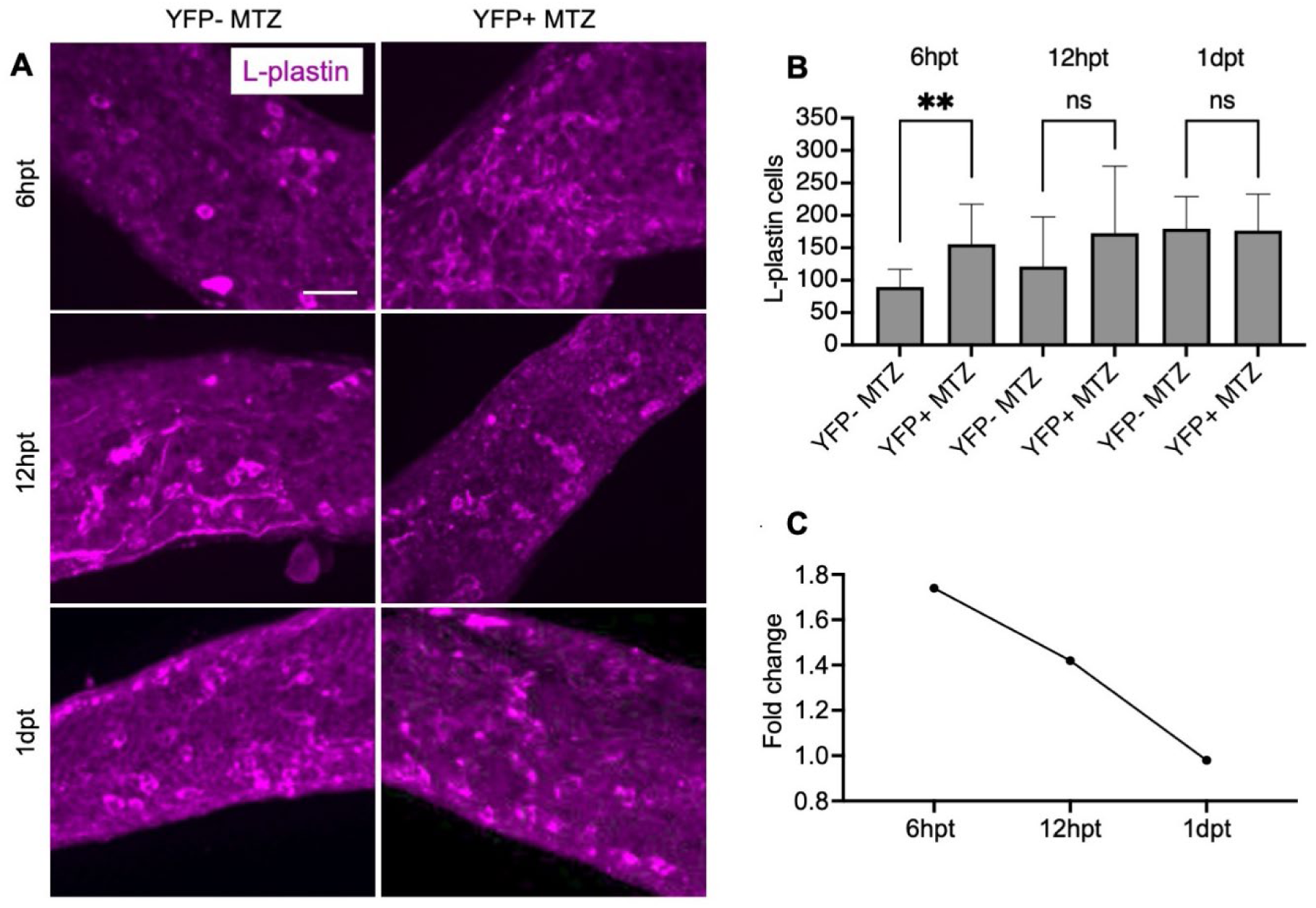
L-plastin+ immune cells increase rapidly after YFP+ cell death in the gut. **(A)** L-plastin+ immune cells (magenta) in YFP-MTZ (non-ablated) and YFP+ MTZ (ablated) larvae at 6 hpt, 12 hpt, and 1 dpt. **(B)** Quantification of L-plastin+ cells at indicated time points. At 6 hpt, YFP+ MTZ groups shows a significant increase in L-plastin+ cells in the gut compared with YFP-MTZ. No significant difference (ns) was detected at 12 hpt and 1 dpt. **(C)** Fold-change analysis of L-plastin+ cell numbers in YFP+ MTZ larvae relative to YFP-MTZ larvae across time points shows a peak at 6 hpt followed by a decline to baseline levels by 1 dpt. **(A)** Maximum projection confocal images of dissected intestines at time point indicated. Data are shown as mean ± SD (n= 10 larvae per group, 2 experiments); Statistical significance: \*\**p* < 0.01; ns = no significant difference; hpt = hours post treatment; dpt = day post treatment; Scale bar = 50 μm.

### YFP+ neurons regenerate following targeted ablation

Next, we investigated whether YFP+ neuron numbers were restored to control levels, which would indicate that they regenerate after MTZ treatment. We quantified YFP+ neurons along the whole gut at different time points before and after treatment in YFP+ MTZ and YFP+ control larvae. YFP+ control larvae exhibited an obvious neuronal network from 0 to 15 dpt (**Figure 5A-B**). In contrast, YFP+ neuronal numbers were reduced in the YFP+ MTZ larvae at 1 dpt, confirming successful ablation of YFP+ neurons (**Figure 5A-B**). By 2 dpt, we find a small number of YFP+ neurons along with developing neurites that bridge across the gut (**Figure S3**). YFP+ neuron numbers continue to increase, but through 6 dpt there was a significant difference in YFP+ numbers compared to the control group. By 9 dpt, regenerated YFP+ neuron numbers reached levels comparable to the controls and remained at control levels through 12 and 15 dpt (**Figure 5A-B**). We noticed that approximately half of the YFP+ MTZ larvae exhibited abnormal body posture after 9 dpt. This phenotype might be caused by off-target ablation of reporter-positive cell in the spinal cord area (**Figure S4**). To investigate whether this affected YFP+ neuron regeneration, we compared regenerated YFP+ neuron counts in larvae with normal posture and those with defective posture at 12 dpt but found no significant difference (**Figure S5**). This indicates that postural defects did not impact neuron regeneration. To determine if there were differences in the timeline of neuron regeneration within discrete gut regions compared to the whole gut, we quantified YFP+ neurons in three main gut regions - anterior, midgut, and posterior (Jianlong Li et al., 2020; Wallace et al., 2005). We found that the timeline of YFP+ neuron regeneration was the same in the three gut regions as in the whole gut (**Figure 5C, B-F**).

**Figure 5.**
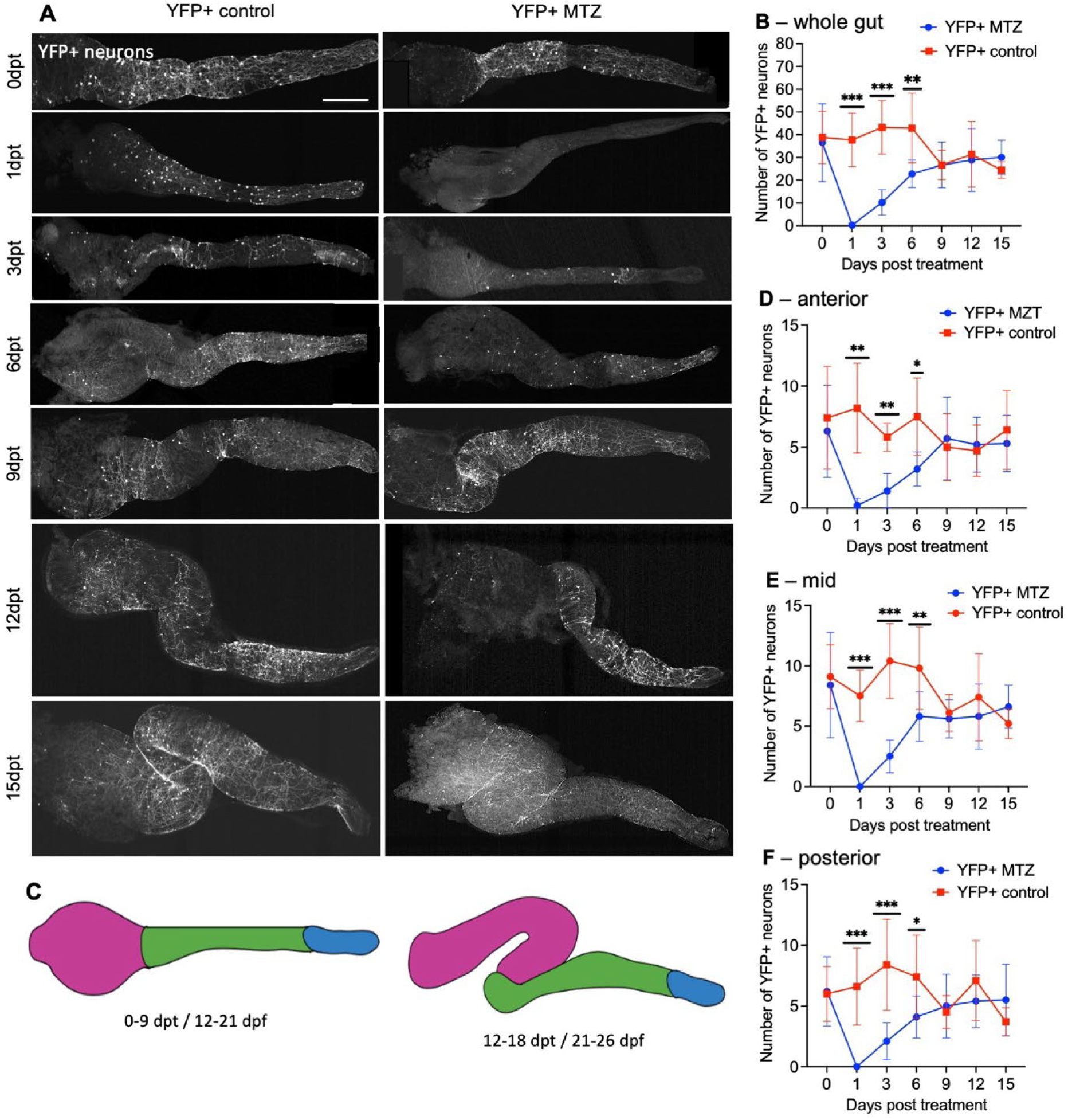
Timeline of YFP+ neuron regeneration in the whole intestine and defined gut regions. **(A)** YFP+ neurons (labelled with anti-GFP antibody) and the neuronal network in YFP+ control and YFP+ MTZ zebrafish larvae at 0 (before treatment) and 1, 3, 6, 9, 12, and 15 dpt. **(B)** YFP+ control larvae exhibited consistent GFP+ neuron numbers till 6 dpt with a slight decrease by 9 through 15 dpt. In contrast, the YFP+ MTZ group showed a dramatic decrease in YFP+ neurons, confirming complete ablation of YFP+ neurons (*p* < 0.001) at 1 dpt. A small number of YFP+ neurons appeared by 3 and 6 dpt, but YFP+ neuron numbers were still significantly decreased compared to the control. By 9 dpt, neuronal number fully recovered with no significant difference between treated and controls at 9, 12 and 15 dpt. **(C)** Schematic representation of the zebrafish gut regions shows the three analyzed gut regions (magenta = anterior, green = midgut, blue = posterior) at 0–9 dpt and 12–18 dpt based on Jianlong Li et al. (2020) and Wallace et al. (2005). Anterior **(D)**, mid **(E)** and posterior **(F)** gut regions show the same timeline of YFP+ neuron regeneration as the whole gut **(B)**. **(A)** Maximum projection confocal images of dissected intestines at time point indicated; Data are presented as mean ± SD (10 larvae per group, 2 experiments); Statistical significance: \*\*\**p* < 0.001, \*\**p* < 0.01 and \**p* < 0.05; dpt = days post treatment; Scale bar = 100 μm.

As shown above, YFP+ neurons colocalized with a small subset of HuC/D+ neurons, indicating that only a fraction of ENS neurons were ablated (**Figure 2D-F**). To assess if ablation of YFP+ neurons has any effects on the YFP-ENS neuronal population, we quantified HuC/D+ neurons in YFP+ MTZ and YFP+ control groups at 0 dpt (before treatment) and 1 and 3 dpt. We found no significant difference in HuC/D+ neuron numbers between treated and control groups after ablation of YFP+ neurons (**Figure S6A-D**). This lack of significance may be due to the relatively small proportion of YFP+ neurons within the total HuC/D+ population—on average, 44.4 out of 204.6 neurons (**Figure 2F**). Additionally, MTZ treatment alone in YFP-MTZ treated larvae did not significantly decrease overall HuC/D numbers (**Figure S6E**).

### Specific co-expressed neuronal subtypes regenerate following ablation

The ENS is composed of various neuron subtypes which control the different functions of the gut including gut motility (Holmberg et al., 2006; Roach et al., 2013; Uyttebroek et al., 2010). The subtypes can be grouped into three main classes which include nitrergic/VIPergic inhibitory motor neurons [neuronal nitric oxide synthase+ (nNOS+); *vasoactive intestinal peptide*+ (*vip*+)], cholinergic excitatory motor neurons [*solute carrier family 18 member 3a*+ *slc18a3a*+)], and serotonergic neurons [5-hydroxytryptamine+, 5-HT+)] (Kuil et al., 2023; Li et al., 2025; Uyttebroek et al., 2010). To determine the neuronal subtype composition within the YFP+ neuronal population, we analyzed co-expression of these subtype markers with YFP at 12 dpf, prior to MTZ treatment. At 0 dpt, we found that 57.54%, 12.26%, 13.33%, and 6.58% of YFP+ neurons co-expressed nNOS, 5-HT, *vip*, and *slc18a3a*, respectively, indicating the presence of distinct YFP+ neuronal subtypes prior to ablation, which fall into the three major classes of ENS neurons [(Kuil et al., 2023; Li et al., 2025; Uyttebroek et al., 2010), **Figure S7**].

To examine if all YFP+ expressing subtypes regenerated, we analyzed the number of YFP+/subtype+ neurons in the MTZ-treated group at 9 dpt and compared them to untreated control and baseline at 0 dpt control. Notably, the number of YFP+/nNOS+ neurons remained unchanged between 0 control and 9 dpt control, and there was no significant difference between MTZ-treated and control group, suggesting robust stability and regenerative capacity for YFP+/nNOS+ neurons (**Figure 6A, B**). Analysis of YFP+/5-HT+ neurons also revealed no significant difference between 0 dpt control and the 9 dpt control group. However, in the MTZ-treated group, the number of YFP+/5-HT+ neurons at 9 dpt remained significantly lower compared to control, indicating a limited regenerative capacity or delayed regenerative kinetics for this neuronal subtype (**Figure 6C, D**).

**Figure 6:**
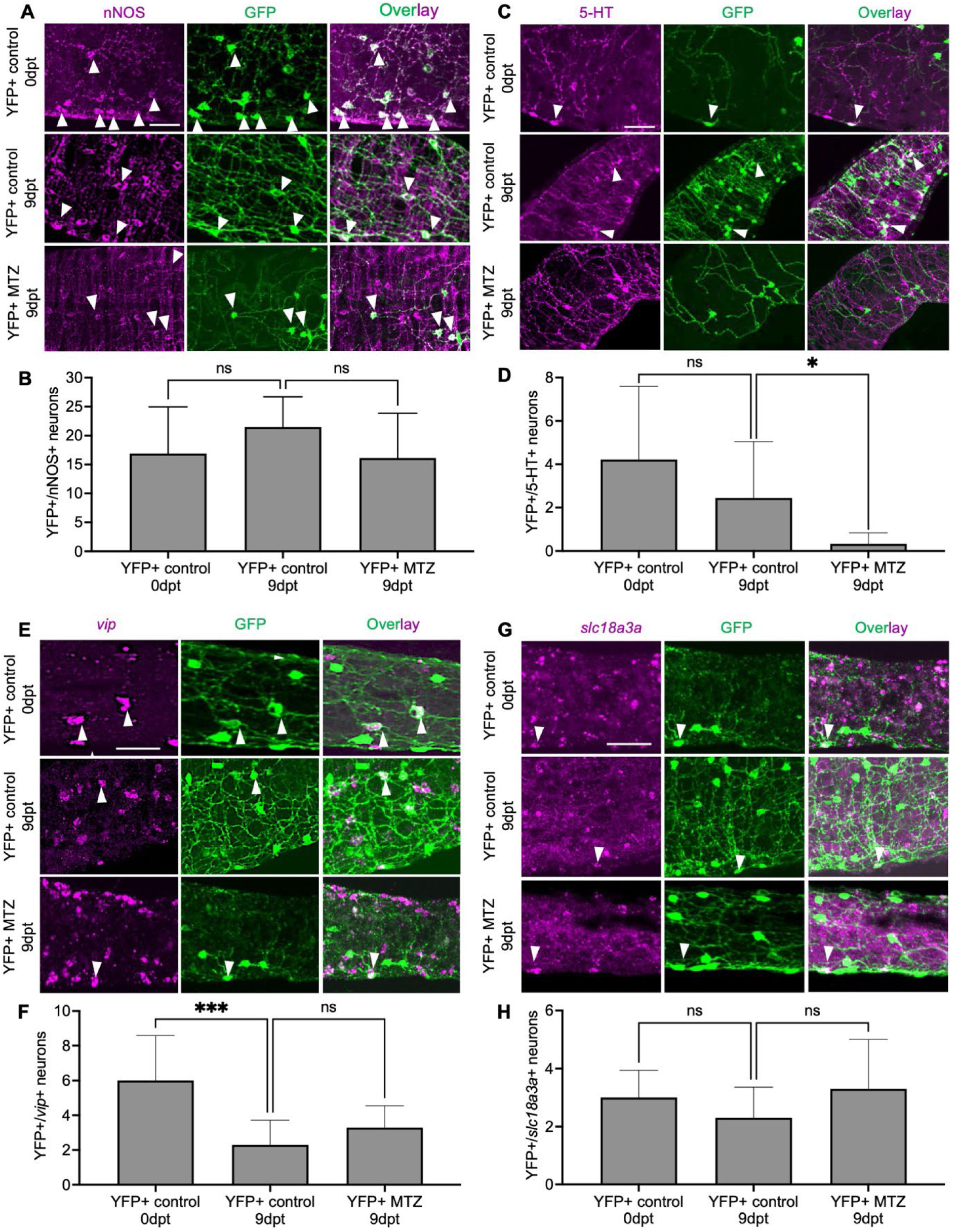
YFP+/nNOS+, YFP+/*vip*+ and YFP+/*slc18a3a*+ but not YFP+/5-HT+ neurons regenerate following ablation. **(A)** Example of nNOS+ (magenta) and YFP+ (labeled with anti-GFP antibody, green) neurons that co-localize (overlay) at 0 dpt control (before treatment) and at 9 dpt in YFP+ control and YFP+ MTZ groups (white arrowheads). **(B)** Quantification of GFP+/nNOS+ neurons at 0 dpt control and 9 dpt in YFP+ control and YFP+ MTZ shows no significant difference. **(C)** Example of 5-HT (magenta) and YFP+ (labeled with anti-GFP antibody, green) neurons that co-localize (overlay) at 0 dpt control and 9 dpt in YFP+ control and YFP+ MTZ groups (white arrowheads). **(D)** Quantification of GFP+/5-HT+ at 0 dpt control and 9 dpt in YFP+ control and YFP+ MTZ. There is no significant difference between 0 dpt and 9 dpt in the YFP+ control group, but GFP+/5-HT+ neurons are significantly reduced in in the MTZ-treated group compared to the control group at 9 dpt. (**E)** Example of *vip* (magenta) and YFP+ (labeled with anti-GFP antibody, green) neurons that co-localize (overlay) at 0 dpt control (before treatment), and 9 dpt in YFP+ control and YFP+ MTZ (white arrowheads). **(F)** Quantification of GFP+/*vip*+ neuron at 0 dpt control and 9 dpt in both YFP+ control and YFP+ MTZ groups. A significant difference is observed between 0 dpt and 9 dpt control groups. However, there is no significant difference between 9 dpt control and MTZ-treated groups. **(G)** Example of *slc18a3a* (magenta) and YFP+ (labeled with anti-GFP antibody, green) neurons that co-localize (overlay) at 0 dpt control and 9 dpt of YFP+ control and YFP+ MTZ (white arrowheads). **(H)** Quantification of GFP+/*slc18a3a*+ neurons shows no significant difference between the groups. **(A, C, E, G)** Maximum projection confocal images of dissected intestines at stage indicated. Data are presented as mean ± SD (n = 10 larvae per group, 2 experiments)_;_ Statistical significance: \*\*\**p* < 0.001 and \**p* < 0.05; ns = non-significant difference; dpt = days post treatment; Scale bars = 10 µm.

Using HCR to detect *vip* and *slc18a3a* expression, we found that the number of YFP+/*vip*+ neurons significantly decreased within the control group between 0 and 9 dpt. However, no significant difference was observed between the MTZ-treated and control groups at 9 dpt, suggesting that YFP+/*vip*+ neurons had successfully regenerated by this time point (**Figure 6E, F)**. Lastly, analysis of the cholinergic marker *slc18a3a* showed no significant differences across groups, indicating that YFP+/*slc18a3a*+ neurons remained stable in the control group and were successfully restored following regeneration (**Figure 6G, H**).

Overall, these findings suggest that the majority of YFP+ cells are nitrergic neurons before ablation. After regeneration, YFP+/nNOS+, YFP+/*vip*+ and YFP+/*slc18a3a*+ neurons are successfully restored. In contrast the YFP+/5-HT+ neurons exhibited lower and/or slower regenerative capacity compared to the other subtypes.

### Decreased gut wave movements recover to control levels after neuronal regeneration

ENS neurons play a critical role in regulating gut motility. Excitatory motor neurons mediate muscle contractions and inhibitory motor neurons induce muscle relaxation (Furness, 2012; Kunze & Furness, 1999). We find that major portion of YFP+ neurons co-express inhibitory and excitatory motor neurons markers (**Figure S6**). Thus, we aimed to determine i) whether the loss of YFP+ neurons affects gut motility and ii) whether the regenerated neurons restore gut function after ablation.

To assess this, we examined one measure of gut motility, the time of the Peristaltic Wave Interval (PWI) of anterograde movements (Holmberg et al., 2006). We measured the PWI at three different time points at 1 dpt, 3 dpt and 9 dpt in YFP+ and YFP-MTZ treated fish, and YFP+ untreated controls. We included the YFP-MTZ treated group to determine if MTZ treatment alone impacted PWI. At 1 dpt, we found no significant difference in the PWI time between control and treated groups (**Figure 7A, Supplementary Videos S1-3**). However, at 3 dpt, the PWI time in the YFP+ MTZ treated group increased significantly compared to both control groups, suggesting slower peristaltic movement in the YFP+ MTZ group (**Figure 7B, supplementary videos S4-6**). At 9 dpt, the PWI time in the YFP+ MTZ treated group was back to control levels (**Figure 7C, supplementary videos S7-9**). Overall, our results suggest that gut motility function is affected after YFP+ neuronal loss and then restored after regeneration, indicating functional recovery.

**Figure 7.**
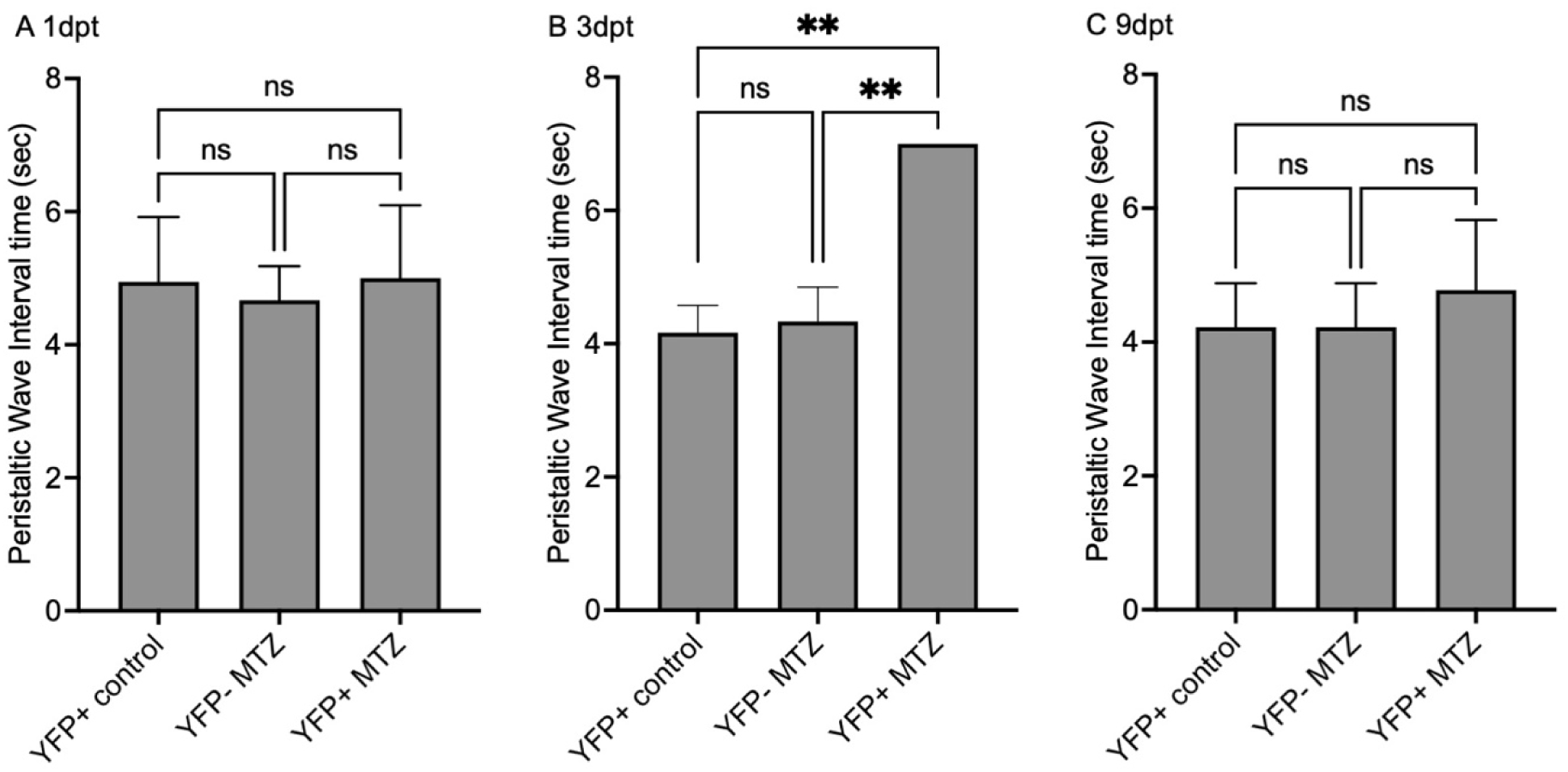
Peristaltic Interval Wave time recovers at 9 dpt after YFP+ neuron ablation. **(A)** At 1dpt, there is no significant difference in gut PWI time across all experimental groups, YFP+ MTZ treated, YFP+ untreated control, YFP-MTZ treated. **(B)** At 3dpt, YFP+MTZ treated larvae showed significantly slower gut PWI time compared to both control groups. **(C)** At 9dpt, there was no significant difference in PWI across all experimental groups. Data are presented as mean ± SD (n = 6 larvae per group, 2 experiments); Statistical significance: \*\**p* < 0.01; ns = non-significant; dpt = days post treatment.

## Discussion

The topic of reparative regeneration has gained significant attention over the past decade due to its potential therapeutic implications, for example to treat ENS diseases with loss of ENS neurons. Yet, to date, no study has demonstrated functional ENS neuron regeneration either in mammalian models or in zebrafish. In the present study, we show targeted chemogenetic ablation and functional regeneration of ENS neurons for the first time in zebrafish (**Figure 8**). Our results determined the spatial and temporal cellular dynamics of targeted neuron ablation as demonstrated by increased TUNEL labeling at 6hpt. Additionally, we observed an acute L-plastin+ immune cell response at 6 hpt that resolves within 1 dpt (**Figure 8**). We find full recovery of ablated neuron numbers by 9 dpt for the majority on ENS neuron subtypes investigated. Specifically, among the regenerated neurons, we found full recovery of nitrergic, cholinergic, and VIPergic subtypes, but only partial or delayed regeneration of serotonergic neurons. Importantly, we also showed functional impacts of neuronal ablation on a gut motility parameter and its functional recovery suggesting functional regeneration in our system (**Figure 8**).

**Figure 8.**
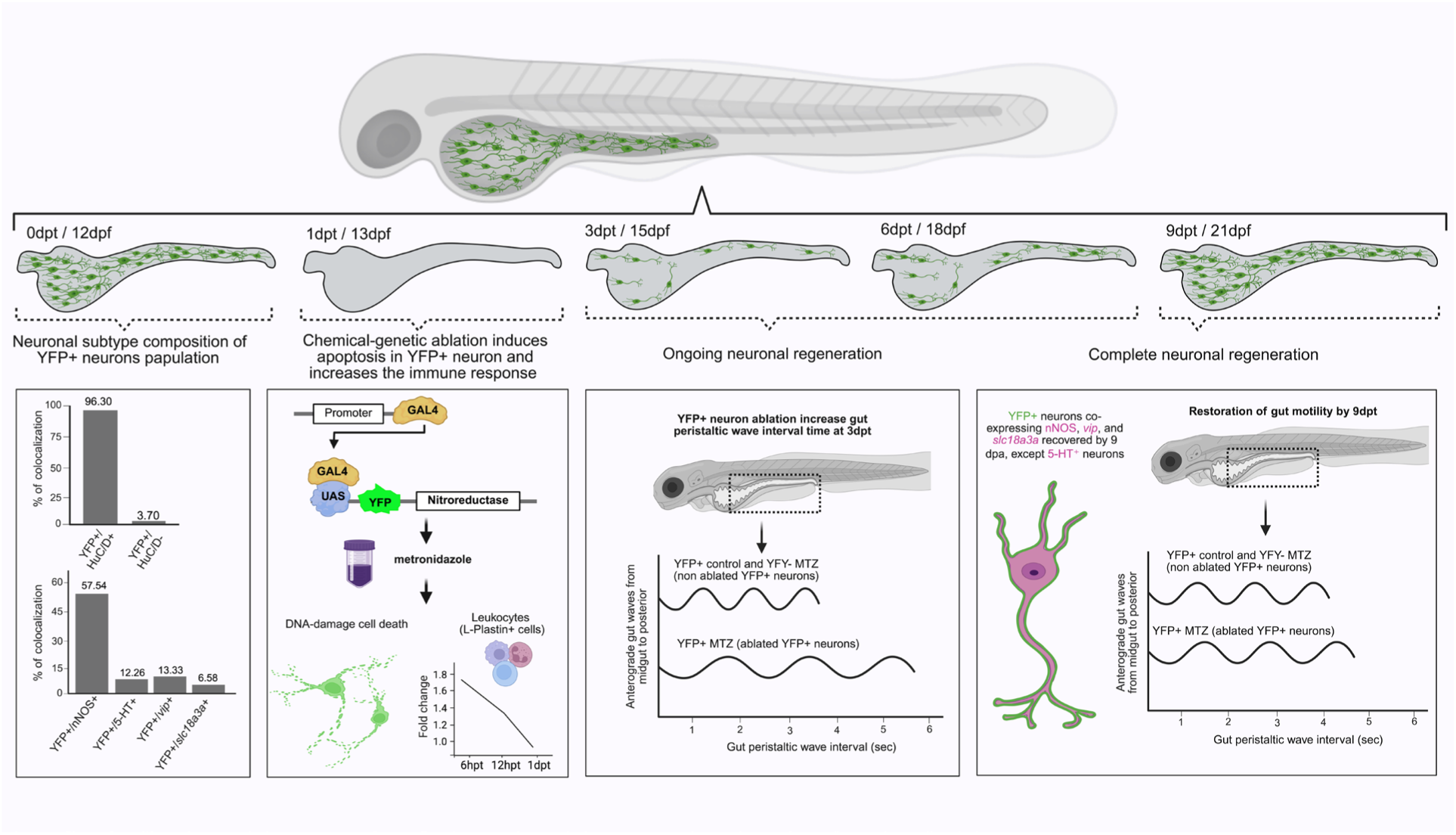
Timeline of events after targeted ablation and regeneration of ENS neurons in zebrafish. Timeline from 0 dpt / 12 dpf to 9 dpt / 21 dpf depicts the different events: i) Neuronal subtype composition of YFP+ neuron population at 0 dpt; ii) Chemogenetic ablation approach activates cell death of YFP+ neurons to completely ablate YFP+ neurons; iii) neuronal regeneration at 3 and 6 dpt which affects gut peristalsis at 3dpt; iv) complete regeneration of YFP+ cells and functional recovery at 9 dpt. **Panel 1** (0dpt / 12dpf): YFP+ cells co-localize with the pan-neuronal marker HuC/D and various neuronal subtypes (nNOS, 5-HT, *vip*, and *slc18a31a*). **Panel 2** (1dpt / 13dpf): Set-up of Gal4-UAS:NTR 2.0 based ablation system. MTZ treatment induces DNA-damage cell death in YFP+ neurons as confirmed by TUNEL assay at 6hpt. Additionally, there is an acute accumulation of L-plastin+ immune cells at 6hpt that resolves within 24 hours. **Panel 3** (3dpt / 15 dpf): Neuronal ablation resulted in increased peristaltic wave interval time, reflecting reduced gut motility. **Panel 4** (9dpt / 21dpf): Complete recovery of YFP+ neurons and most neuronal subtypes (nNOS+, *vip*+, *slc18a31*+), except 5-HT+ neurons and full functional recovery of gut motility. dpf = days post fertilization, dpt = days post treatment.

### Extensive ENS neuron ablation and their complete regeneration in zebrafish

In zebrafish, the only two studies to study ENS neurons regeneration used focal laser ablation to target a few ENS neurons in a small gut region. Both studies reported generation of new neurons following ablation (El-Nachef & Bronner, 2020, Ohno et al., 2021). However, quantification of regenerated neurons showed that they did not return to control levels in the analyzed time frame, nor were impacts on gut function investigated (Ohno et al., 2021). In our study, using chemogenetic ablation of ∼44.4 ENS neurons, we find robust regeneration of ablated cells to control levels by 9 dpt which stay at control levels through 15 dpt, the last time point that we analyzed. We find that different neuronal subtypes within the three main ENS neuron classes are able to regenerate but identify interesting subtype-specific differences in regenerative capacity or timing (**Figure 8**). Our work also shows for the first time recovery of a parameter of gut function - gut wave propagation - indicative of successful functional regeneration (**Figure 8**). In addition, our approach has several experimental advantages compared to laser ablation. First, laser ablation is less efficient, as it can only ablate a limited number of neurons within specific gut regions (El-Nachef & Bronner, 2020, Ohno et al., 2021). For instance, one study only ablated ∼16 neurons within one-somite-width of the posterior gut at 10-15 dpf (Ohno et al., 2021). In contrast, our chemogenetic ablation system successfully ablated on average 44 neurons throughout the whole gut, significantly improving efficiency. Second, laser ablation is time-consuming as it requires individual larval preparation and precise manual targeting (Ohno et al., 2021). In contrast, our methods enable the simultaneous ablation of ENS neurons in several hundred of larvae. Third, laser irradiation can cause unintended damage to non-neuronal, surrounding tissues, including smooth muscle and epithelial cells, which may impact regeneration and complicate downstream analyses such as gut motility. In contrast, our approach is spatiotemporally controlled ablation that precisely targets only specific ENS neurons and had no deleterious effects on the overall neuronal population in the gut (**Figure 8**).

### Zebrafish are a powerful model system to study ENS regeneration

So far, ENS regeneration research has mostly focused on mammalian model systems with only limited work in non-mammalian models such as zebrafish (Corvetti et al., 2001; El-Nachef & Bronner, 2020; Gabella & Trigg, 1984; Galligan et al., 1989; Hanani et al., 2003; Jew et al., 1989; Joseph et al., 2011; Karaosmanoǧlu et al., 1996; Laranjeira et al., 2011; Luck et al., 1993; Ohno et al., 2021; Stavely et al., 2024; Tokui et al., 1994). Previous studies in mammalian models involving surgical (Galligan et al., 1989; Karaosmanoǧlu et al., 1996; Tokui et al., 1994), stenosis (Corvetti et al., 2001; Gabella & Trigg, 1984; Jew et al., 1989; Joseph et al., 2011), and chemical (Hanani et al., 2003; Laranjeira et al., 2011; Luck et al., 1993) ablation methods consistently report incomplete neuronal regeneration, and gut region- or species-specific differences in regenerative outcomes of ENS neurons (Rueckert and Ganz, 2022). Notably, all previous approaches also reported structural abnormalities at the injury site by co-lateral damage of surrounding tissues. To address this, recent works in adult mice reported focal ablation specifically of ENS neurons using a diphtheria toxin-based chemogenetic ablation approach (Bhave et al., 2019; Stavely et al., 2024). Even though this study reported that mature, post-mitotic enteric neurons in adult mice can regenerate axons and reinnervate the gut after injury, there was still limited regeneration of neurons even three months after ablation (Stavely et al., 2024). This indicates that mammals have generally limited ENS regenerative abilities. In contrast, the present study demonstrates that zebrafish are a powerful model to study the signals and cellular-molecular processes underlying functional ENS regeneration. Our spatio-temporally controlled approach ablates approximately one fifth of all ENS neurons by inducing cell death with subsequent full recovery after only 9 days. Ablation along the entire gut will enable the analysis of cellular responses and gene regulatory programs active during ENS regeneration as whole guts can be analyzed. We find an acute immune response during the time of cell death which normalizes at 1 dpt. Studies on adult zebrafish brain traumatic injury and selective rod photoreceptor ablation in the retina have demonstrated that immune responses are crucial components of regenerative processes (Kyritsis et al., 2012; White et al., 2017). In our study, selective ablation of enteric neurons (**Figure 8**) led to a marked accumulation of L-plastin+ cells as early as 6 hours after MTZ treatment. Given that L-plastin is strongly expressed in leukocytes, including macrophages, monocytes, and neutrophils (Hiroto Shinomiya, 2012), this finding suggests that selective neuronal death in the ENS triggers an acute immune cell recruitment, similar to what has been described in models of traumatic brain injury and retinal regeneration (Kyritsis et al., 2012; White et al., 2017). Future studies will be needed to provide mechanistic insights into how immune–neural interactions (Graves et al., 2021) regulate ENS regeneration. Our approach thus provides a rapid experimental setup to identify the signals and cellular responses important for neuronal regeneration in the ENS in the future (**Figure 8**).

### Regeneration of neurons in the zebrafish CNS *vs* ENS

The zebrafish central nervous system has been extensively studied with regards to its regenerative capacity (Kaslin et al., 2009, 2013; März et al., 2010) while little is known about ENS regeneration (Ohno et al., 2021). In the central nervous system, brain regions show different capacities to regenerate neuronal subtypes. In the zebrafish telencephalon, all ablated neuron types regenerate (Barbosa et al., 2015; Kroehne et al., 2011; Reimer et al., 2008), whereas the cerebellum is capable of regenerating many, but not all neuron types (Kaslin et al., 2017). This variable regenerative capacity of the cerebellum has been attributed to varying ability of cerebellar stem and progenitor cells to generate neurons and glia throughout the lifespan (Kaslin et al., 2009, 2013). We also find differences in regeneration of neuron types in the ENS where some subtypes (nitrergic, cholinergic and VIPergic) regenerate fully, whereas serotonergic neurons do not reach control levels (**Figure 8**). There could be three possibilities for this difference in regeneration: (1) serotonergic neurons take longer to regenerate, (2) the progenitor cells or cellular mechanisms to regenerate serotonergic neurons are not activated at this stage of post-embryonic development, and (3) the low level of serotonergic cell loss incurred is insufficient to trigger serotonergic cell replacement. Further work will determine which cellular responses are triggered after cell ablation and the progenitor population(s) that generate new ENS neurons and neuronal subtypes in zebrafish.

## Supplementary Materials

The following supporting information can be downloaded at: https:

## Author Contributions

Conceptualization: M.A.S, K.M., H.R., J.G. Methodology, Data curation, Validation, Formal Analysis and Investigation: M.A.S, K.M., H.R. Visualization: M.A.S. Writing—Original Draft Preparation: M.A.S. Writing—Review and Editing: M.A.S, K.M., H.R., A.V.S., D.F.A., J.S.M., and J.G. Supervision, Project Administration, Funding Acquisition and Recourses: D.F.A., J.S.M., J.G. All authors have read and agreed to the published version of the manuscript.

## Institutional Review Board Statement

The animal study protocol was approved by the Michigan State University Institutional Animal Care and Use Committee.

## Acknowledgments

We want to thank Rachel Alcorn, Taylor Lawrence, Giovanna Haddock, Kristen Lounsbury, Anthony Turner, Samantha Lampman, and Sharon An for excellent fish care.

## List of Abbreviations

ENS: Enteric nervous system
MTZ: Metronidazole
HSCR: Hirschsprung Disease
DPF: Days post fertilization
DPT: Days post treatment
HPT: Hours post treatment
NTR: Nitroreductase
YFP: Yellow fluorescent protein
GFP: Green fluorescent protein
UAS: Upstream activating sequence
PWI: Peristaltic Wave Interval
TdT: Terminal deoxynucleotidyl Transferase
BSA: Bovin serum albumin
PBS: Phosphate-Buffered Saline
HCR: Hybridization Chain Reaction
SD: standard deviation

## Supplementary figures

**Figure S1:**
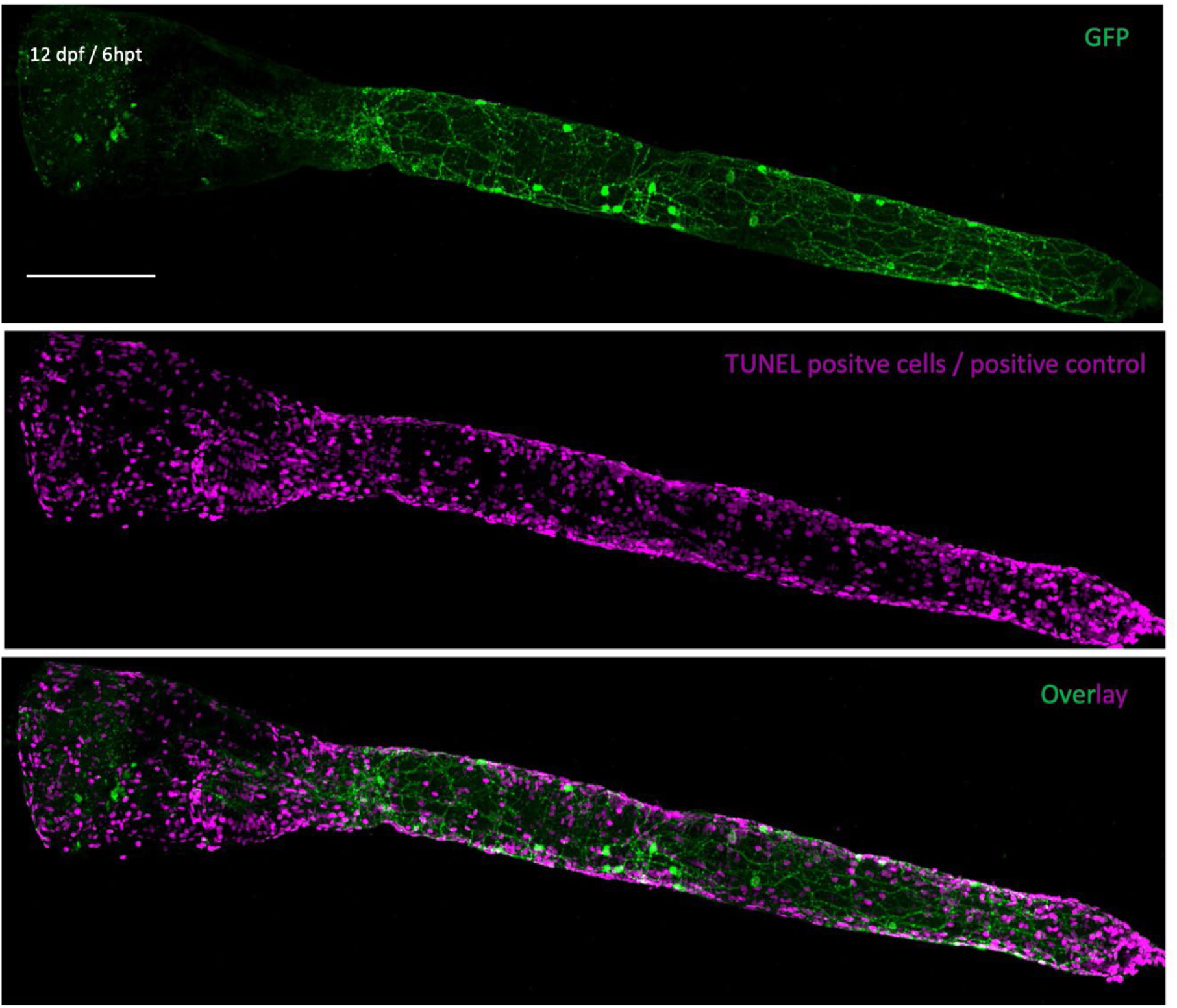
Inducing cell death shows an expected increase in TUNEL+ cells. Inducing cell death as a positive control results in a high number of TUNEL-positive cells (magenta, middle panel) in YFP+ control (green) larvae at 6 hpt (n= 9 larvae, 2 experiments). Maximum projection confocal images of dissected intestine at stage indicated. Scale bar = 100 μm; hpt = hours post treatment.

**Figure S2.**
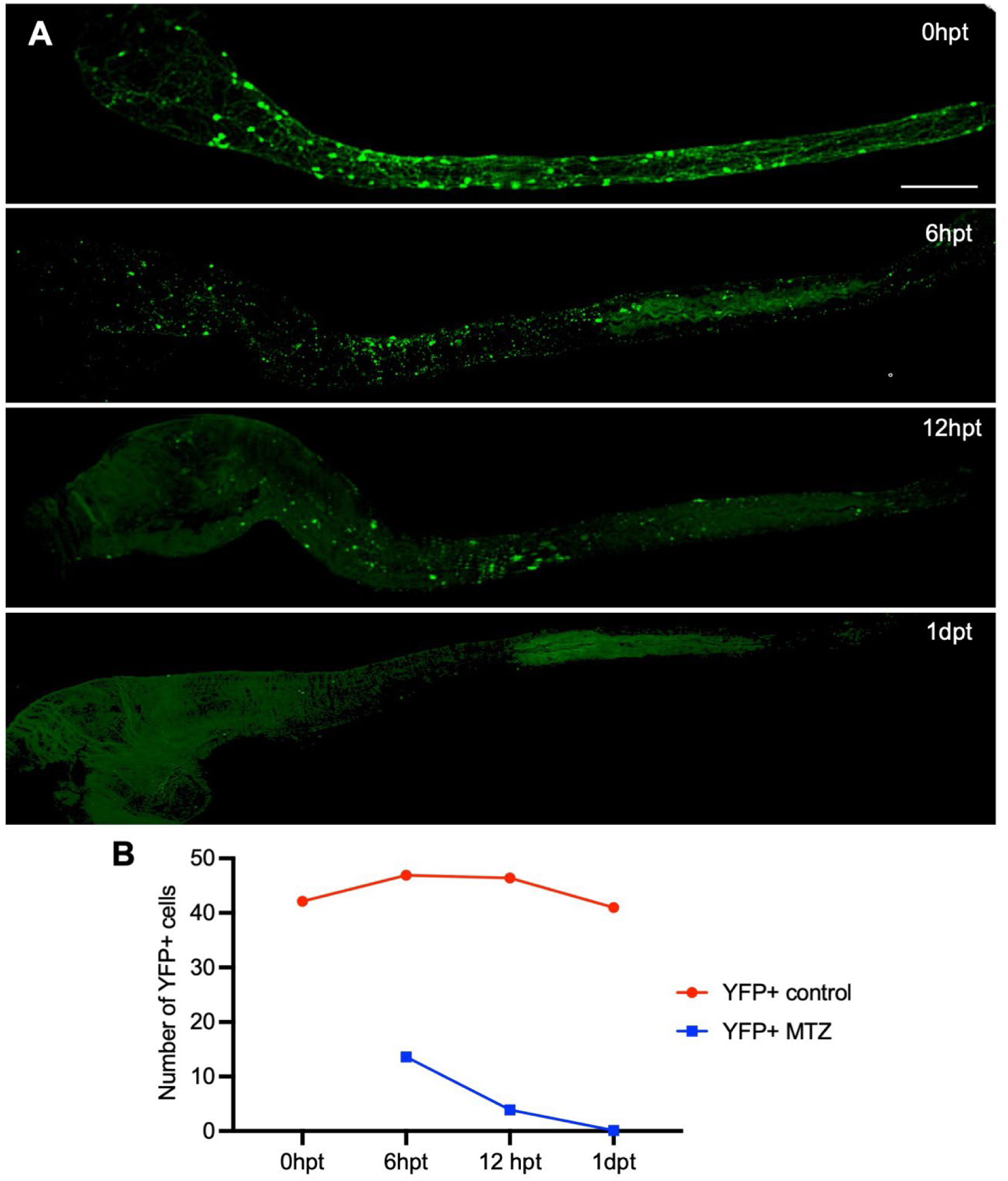
Metronidazole (MTZ)-induced ablation of YFP+ neurons is completed within 24 hours post-treatment. **(A)** YFP+ neurons (labeled with anti-GFP, green) are present at 0 dpt (before treatment of MTZ), but YFP+ neuron numbers continuously decrease after treatment of MTZ at 6 hpt and 12 hpt, and are completely gone at 1 dpt. **(B)** Quantification of YFP+ neurons at time points indicated in YFP+ control (red line) and YFP+ MTZ groups (blue line). The YFP+ control group maintains stable cell numbers across all time points. In the YFP+ MTZ-treated group YFP+ neuron numbers rapidly decline at 6 hpt, with near-complete ablation by 1 dpt. **(A-C)** Maximum projection confocal images of dissected intestines at time point indicated. Data presented as mean (n = 10 larvae per group, 2 experiments); hpt = hours post-treatment, dpt = hours/days post-treatment; Scale bar = 100 μm.

**Figure S3.**
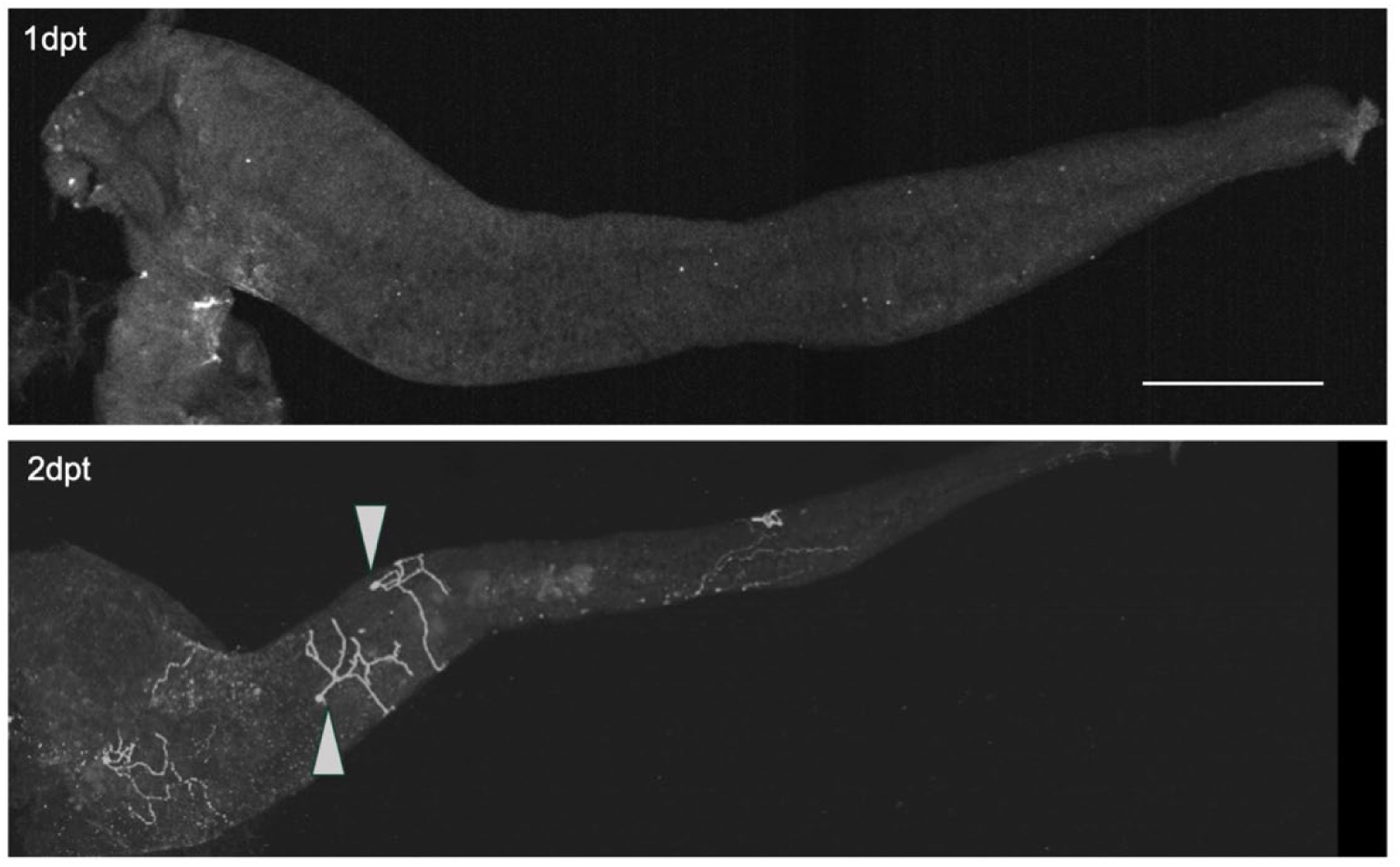
Regenerating neurites in YFP+ neurons following ablation. At 2 dpt, newly generated YFP+ neurons form neurites as the beginning step of neuron regeneration. Upper panel: at 1 dpt there is complete ablation of YFP+ neurons. Lower panel: at 2 dpt developing neurites that form a bridge (arrow) between the newly generated neurons (n = 6 larvae per group. 2 experiments). Maximum projection of confocal image of dissected intestine at time point indicated. dpt = days post treatment; Scale bar = 100 μm.

**Figure S4.**
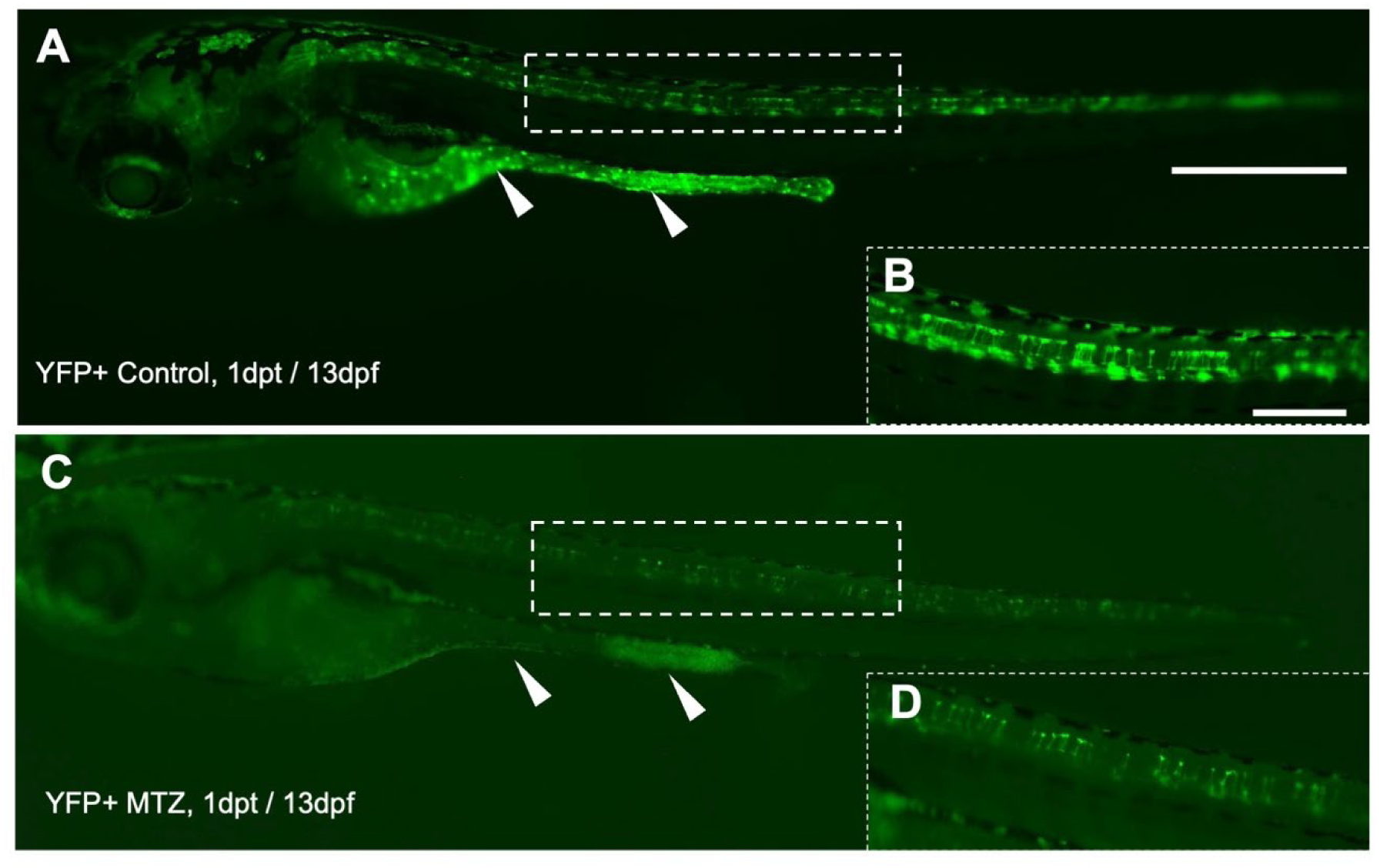
Off-target ablation of YFP+ cells from the spinal cord region. **(A)** In YFP+ control larvae, the intestine contains a network of YFP+ ENS neurons along the gut and in cells within the area of the spinal cord and in the brain. **(B)** Magnified view shows YFP+ cells in the spinal cord area. **(C)** In YFP+ MTZ treated larvae, YFP+ ENS neurons are essentially absent. **(D)** Magnified view shows that YFP+ cells in spinal cord area are also reduced after MTZ treatment, demonstrating off-target NTR-mediated cell ablation in the spinal cord. (**A-C)** Whole-mount side views of live images of YFP+ control (untreated) and YFP+ MTZ (treated) at stages indicated. Arrowheads indicate reporter-positive cells in the gut region. Dashed box outlines the region for magnified view. dpt = days post treatments; (A, C) scale bar = 100 μm; (B, D) scale bar = 20 μm.

**Figure S5.**
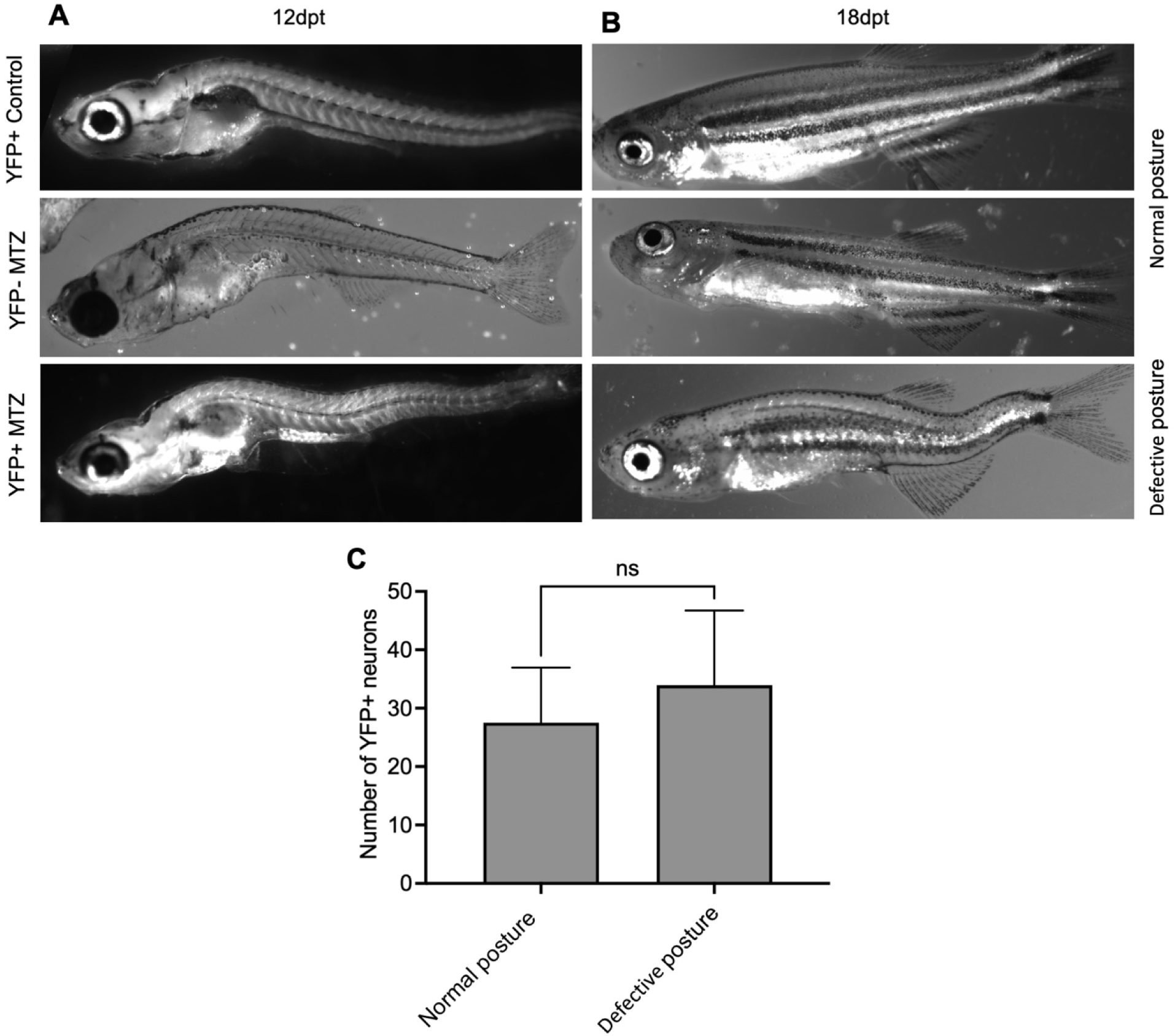
Abnormalities in body posture do not affect YFP+ neuron regeneration. **(A-B)** Whole-mount side views of larvae at stage indicated from treated (YFP-/+ MTZ treated) and Control (YFP+ untreated) groups. Only half of YFP+MTZ treated shows the body posture defect after 9 dpt. **(C)** The bar graph shows the number of regenerated YFP+ neurons in normally developed larvae compared to those with defective body posture at 12 days post treatment. Data presented as mean ± SD (n = 8 larvae per group, 2 experiments); ns = no statistically significant differences; Scale bar = 100 µm.

**Figure S6.**
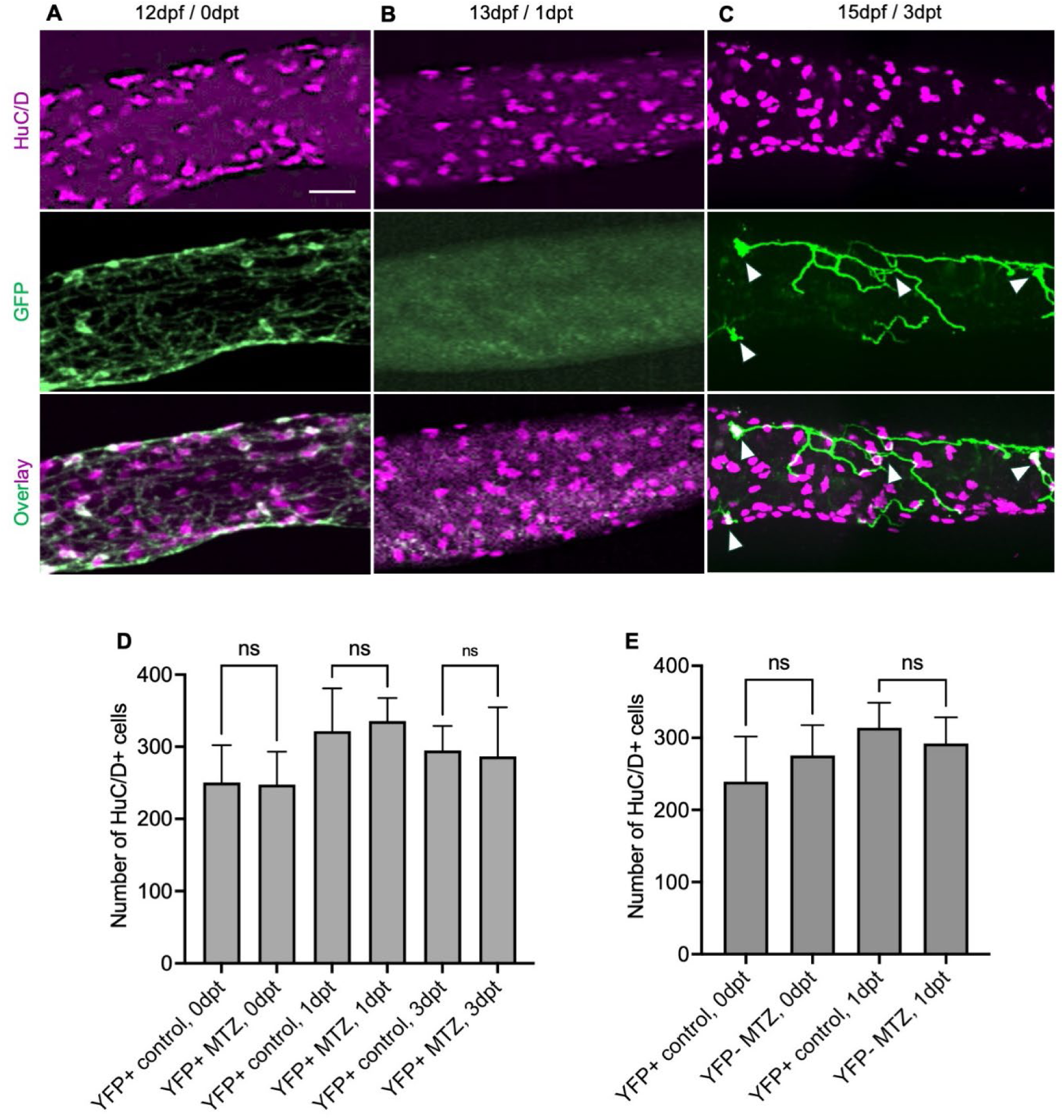
YFP+ neuron loss and MTZ treatment of YFP-larvae does not affect neuron numbers. **(A)** At 12 dpf (before treatment, 0 dpt), the pan-neuronal marker HuC/D (magenta) and YFP+ cells (labeled with anti-GFP, green) show high colocalization (overlay). **(B)** Following MTZ treatment at 13dpf (1dpt), GFP-HuC/D+ cells are present, whereas GFP+ cells are completely absent due to ablation of YFP+ neurons. **(C)** 15dpf (3dpt) shows GFP+ cells and exons (indicated by while arrow) regeneration process. **(D)** Quantification of HuC/D-positive neuron counts before (0 dpt) and after MTZ treatment (1 and 3dpt). There was no significant difference in total HuC/D+ cells between control and treatment group (n = 10 larvae per group, 2 experiments). **(E)** Quantification of HuC/D+ neurons in the gut in YFP-MTZ treated and YFP+ untreated controls at 12dpf (before MTZ treatment) and 13dpf (after MTZ treatment). Bar graph shows the number of HuC/D+ cells. No significant difference was observed between treatment groups (n = 10 larvae, 2 experiments). Data presented as mean ± standard deviation; ns = not significant, dpt = days post treatment. **(A-C)** Maximum projection confocal images of dissected intestines. Scale bar = 100 μm; Data are presented as mean ± SD, ns = not significant; dpf = days post fertilization; dpt = days post treatment.

**Figure S7.**
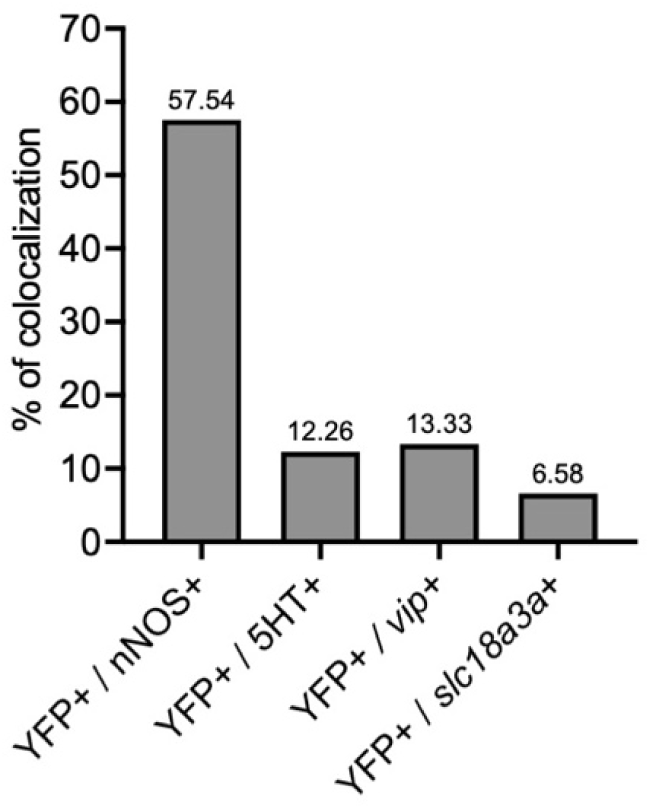
The YFP+ neuronal population consists of different neuronal subtypes at 12 days post fertilization. Quantification of YFP and neuronal subtype colocalization. Bar graph shows the percentage of co-expression of each subtype marker with YFP at 12 days post fertilization / 0 day post treatment. Data presented as mean (n = 10 larvae per group, 2 experiments).

Supplementary videos have been uploaded to the designated repository.

| Groups | Video description |
| --- | --- |
| 1dpt | Video S1 = YFP- MTZ<br>Video S2 = YFP+ control<br>Video S3 = YFP+ MTZ |
| 3dpt | Video S4 = YFP- MTZ |
|  | Video S5 = YFP+ control<br>Video S6 = YFP+ MTZ |
| 9dpt | Video S7 = YFP- MTZ<br>Video S8 = YFP+ control<br>Video S9 = YFP+ MTZ |
| dpt = days post treatment |  |

